# Mechanical forces drive mitochondrial matrix extrusion and apoptotic pore growth

**DOI:** 10.1101/2025.05.12.653510

**Authors:** Andreas Jenner, Timo Dellmann, Hyuntae Kim, David Gomez, Cristiana Zollo, Jürgen Köfinger, Katrin Seidel, Felix Gaedke, Astrid Schauss, Gerhard Hummer, Ana J. Garcia-Saez

## Abstract

Apoptotic pore opening by BAX and BAK at the mitochondrial outer membrane is a key step in the cell commitment to death. The subsequent inner membrane extrusion and permeabilization releases mitochondrial DNA into the cytosol, which can trigger inflammatory signaling. However, the underlying mechanisms have not been elucidated. Here we developed CLOSE microscopy, a multi-correlative approach that enabled the simultaneous analysis of BAK stoichiometry and nanoscale organization in individual apoptotic pore complexes in relation to mitochondrial ultrastructure. We find that the outer membrane opening at the apoptotic pore defines the spatial arrangement of BAK nanoassemblies. We identify mechanical stress as a driver of inner membrane extrusion, which can be perturbed by osmoregulation. The extruded inner membrane in turn exerts forces on the outer membrane that promote apoptotic pore growth, in line with membrane dynamics simulations. Our study reveals a tight interplay between the inner and outer membranes during mitochondrial permeabilization in apoptosis and establishes a biophysical mechanism for inner membrane extrusion that defines the structural organization of the apoptotic pore.

## Introduction

BAX and BAK are proapoptotic members of the BCL-2 protein family and are critical for apoptosis execution ^1,2^. They mediate mitochondrial outer membrane (MOM) permeabilization (MOMP) during apoptosis, a pivotal point in the cell’s commitment to death. MOMP enables the release of apoptotic factors, including cytochrome *c* (cyt *c*) and Smac/DIABLO, from the mitochondrial intermembrane space into the cytosol, initiating the downstream apoptotic caspase cascade ^3–6^. This ultimately results in the dismantling of the cell into apoptotic bodies, which are engulfed by macrophages, usually ensuring an immunologically “silent” removal of the dead cell ^7,8^.

The dynamic oligomerization of BAX and BAK molecules into high-order complexes of heterogeneous size and shape results in the opening of apoptotic pores at the MOM that continue to grow during apoptosis progression. At the nanoscale, these assemblies form line, arc, and ring structures, with arcs and rings capable of forming pores in the membrane ^9–12^. BAX and BAK membrane pores are lined by both proteins and lipids, so that the biophysical properties of the membrane govern the pore size and stability ^13–17^. *In vitro*, BAX membrane insertion has been associated with a reduction in line tension at the pore edge that stabilizes the open pore ^18,19^.

As apoptotic pores form in the MOM, mitochondria undergo functional and structural rearrangements ^20^. The mitochondrial inner membrane (MIM) remodels, including *cristae* unfolding and reorganization ^21–23^. A striking event is the permeabilization and extrusion of the MIM through BAX and BAK apoptotic pores in the MOM ^24,25^. MIM permeabilization (MIMP) releases immunogenic mitochondrial factors, such as mitochondrial DNA (mtDNA), into the cytosol, which can trigger inflammatory responses under conditions of low caspase activity ^26,27^. The growth rate of the apoptotic pore is tuned by the dynamic co-assembly of BAX and BAK, as well as by lipid composition, and regulates the sequential release of mitochondrial contents into the cytosol, impacting inflammatory signaling ^9,12^. As a result, MIMP following MOMP can make apoptosis inflammatory, even under sublethal conditions, as recently reported in models of cellular senescence or tumor cell death ^28–30^.

However, the mechanisms of MIM extrusion and its connection to the nanoscale organization of BAX and BAK at the apoptotic pore in the MOM remain elusive. Addressing this question requires exploration of the discrete assemblies formed by BAX and BAK with regards to their copy number, spatial arrangement and position at organelles subdomains, which is technically challenging. To this aim, we report here a novel multi-correlative microscopy approach that allows the simultaneous investigation of the oligomeric state and supramolecular organization of BAK pores, in relation to the mitochondrial ultrastructural alterations and membrane integrity, at the single-pore level. We show that the oligomeric state is not determinant of the shape of BAK nanoassemblies, which instead delineate MOM openings. The size of MOM openings also impacts MIM extrusion, which follows mechanical forces, as supported by membrane dynamics simulations. Accordingly, MIM extrusion responds to osmotic swelling of the matrix. We find that MIM extrusion also contributes to apoptotic pore growth on the MOM, and thus to its structural arrangement, by exerting forces on its edge, which we quantify. Our findings thus uncover that MIM extrusion and the growth of the apoptotic pore at the MOM during apoptosis are tightly interconnected and establish a biophysical mechanism underlying these processes.

## Results

### CLOSE microscopy, a multi-correlative microscopy approach

To study how BAK oligomerization is related to the supramolecular arrangement of the apoptotic pore and how this affects mitochondrial integrity and ultrastructure, we developed a multi-correlative approach that we named <u>C</u>orre<u>L</u>ative <u>O</u>ligomerization <u>S</u>TED and <u>E</u>lectron (CLOSE) microscopy. This method enables the correlative assessment of the protein stoichiometry of single particles using photon-counting confocal microscopy and their nanoscale organization based on STED microscopy, as well as the cellular context information provided by an additional correlative step with electron tomography (ET) (Figure 1). CLOSE microscopy can be applied to discrete protein complexes in the cell, which we illustrate here by using it to dissect how BAK oligomerization and supramolecular assembly in the apoptotic pore relates to MOMP and MIM extrusion.

**Figure 1:**
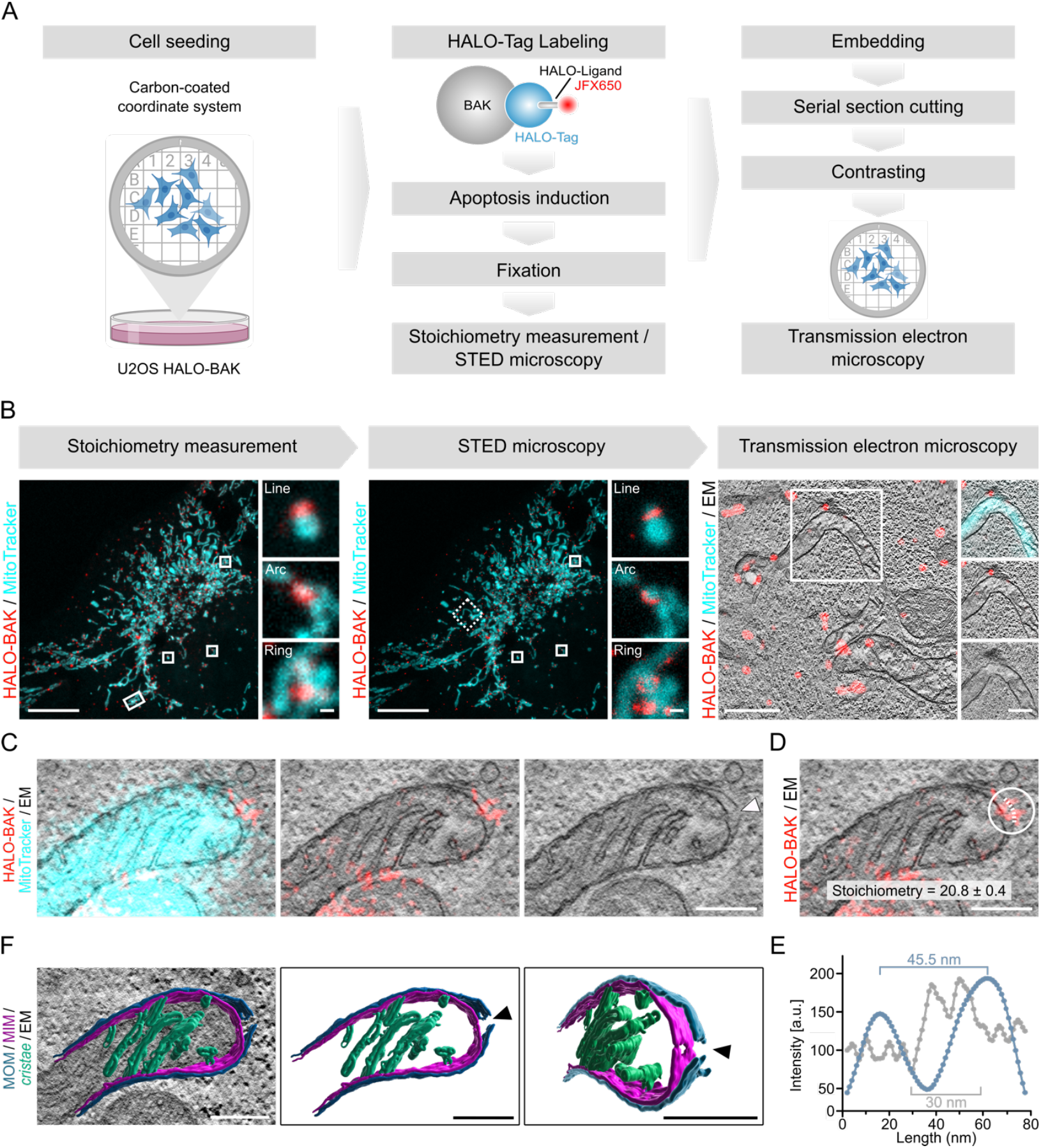
Development of CLOSE microscopy and application to the apoptotic pore. A) Schematic of the workflow including cell seeding on carbon-coated coordinate system gridded dishes, labeling of HALO-BAK with JFX650 HALO-Tag ligand, induction of apoptosis and cell fixation. The stoichiometry of apoptotic BAK is then measured by photon-counting confocal microscopy, followed by STED microscopy to examine the nanoscale structural organization of BAK. Samples are resin-embedded, cut and contrasted for transmission electron microscopy to visualize mitochondrial ultrastructure. B) Representative photon-counting confocal (left), STED (middle), and transmission electron microscopy (right) images of U2OS HALO-BAK cells labeled with JFX650 HALO-Tag ligand (red) one hour after apoptosis induction. Mitochondria were visualized using MitoTracker orange (cyan). Enlarged images correspond to cropped regions of the overview image as indicated, showing representative photon-counting confocal and STED images of line, arc, and ring structures. Dashed rectangle indicates region shown in transmission electron microscopy image. Scale bar 10 µm (zoomed images 500 nm) for stoichiometry and STED images and 1 µm (zoomed images 500 nm) for transmission electron microscopy images. Data are representative of n = 8 independent experiments. C) Overlaid STED-CLEM image of a single mitochondrion showing MitoTracker (cyan, confocal) and BAK (red, STED) fluorescence signal and EM image (grayscale, left panel), BAK signal and EM image (middle panel), and single EM image (right panel). Arrowhead indicates position of MOM discontinuity. Images correspond to the magnified (rotated and flipped) region of the image in (B, left panel) as indicated by the rectangle. Scale bar 250 nm. D) STED-CLEM overlay image of BAK (red, STED) and EM signal (grayscale) of the mitochondrion shown in (C). White circle indicates region of interest for quantification of BAK stoichiometry (as indicated) from photon-counting confocal microscopy image (not shown). Dashed line indicates the region of BAK fluorescence and EM intensity measurement (shown in E) along the MOM discontinuity. Scale bar 250 nm. E) Quantification of BAK fluorescence (blue) and EM intensity (gray) at the line profile indicated by the dashed line in (D). F) 3D rendered MOM (blue), MIM (purple), and *cristae* membranes (green) overlaid with EM image (grayscale, right panel) in top view (middle panel) and tilted view (right panel) of the image shown in (C). Arrowheads indicate the position of the MOM and MIM discontinuities. Scale bar 250 nm.

We seeded cells endogenously expressing HALO-BAK (Figure S1) on a coverslip coated with a coordinate system, and labeled HALO-BAK with the JFX650 HALO-tag ligand and mitochondria with MitoTracker Orange. We induced apoptosis with a combination treatment of BH3-mimetics (ABT-737 and S63845) and fixed the cells at 1 h, shortly after widespread MOMP in the cell population under our experimental conditions, which we estimated using TMRE fluorescence decline due to mitochondrial membrane potential loss as a proxy (Figure S1E-F) ^9,11^. Next, the cells were subjected to photon-counting confocal imaging ^11^ followed by acquisition of a z-stack of the same cells with STED microscopy, to assess the stoichiometry and nanoscale structure of individual apoptotic pores, respectively. This order is important to minimize the impact of photobleaching during STED imaging for stoichiometry assessment by photon counting. The samples were then prepared for transmission ET to measure mitochondrial ultrastructure (Figure 1A). We used the cell position on the coordinate system, which was transferred to the resin like a stamp, to identify the same individual cell across the images and roughly overlay light and electron microscopy images. For fine correlation of the subcellular structures, aligning the mitochondrial network distribution across the images as an internal reference provided excellent results (Figure 1B).

We correlated distinct HALO-BAK nanoassemblies from the STED images with their fluorescence signal in photon counting confocal image. We quantified the copy number of BAK in these individual structures by ratiometric stoichiometry analysis based on the photon counting data as in ^9,31–33^ and using the endogenously expressed 32-mer nuclear pore complex component 96 (NUP96) tagged with HALO (NUP96-HALO) as a calibration standard ^34^ (Figure S2A-B). Next, we correlated the stoichiometry and supramolecular organization of single apoptotic HALO-BAK assemblies with the mitochondrial ultrastructure by overlaying the corresponding STED and ET plane images (Figure 1C). This allowed the examination of the structural integrity of mitochondrial membranes directly at the sites of apoptotic BAK assemblies measured by STED microscopy in the electron tomograms. Figure 1D exemplifies the outcome of this process by identifying a ring-like BAK structure, which we estimated to contain about 20.8 ± 0.4 HALO-BAK molecules and which overlapped with a pore in the MOM. The fluorescence intensity of HALO-BAK along the MOM opening presented a drop inside the HALO-BAK ring, indicating an inner pore diameter of *∼*45 nm, which overlapped with the MOM discontinuity of *∼*30 nm, as measured in the EM tomogram (Figure 1E).

Interestingly, in some cases we detected MIM discontinuities indicative of MIMP directly at sites of HALO-BAK assemblies associated with MOM pores (Figures 1C and S3). Segmentation of the mitochondrial compartments in the tomogram clearly showed a pore at the MIM underneath the MOM HALO-BAK pore (Figures 1F and S3D-E; Supplementary Movie 1). The visualization of a MIM pore that correlates with apoptotic pores at the MOM with the MIM contained within the MOM reveals that MIMP can happen prior to extrusion. However, while the distribution of HALO-BAK at MOM openings suggests that it does not participate in MIMP (see below), we cannot draw definitive conclusions from our data regarding the mechanism of MIM pores.

### The spatial arrangement of BAK nanoassemblies is not determined by their stoichiometry

We then performed a statistical analysis of the stoichiometry of HALO-BAK assemblies relative to their nanoscale organization. Consistent with previous studies ^9^, we found a broad distribution of HALO-BAK oligomeric states of individual apoptotic pores, ranging from 50 to over 500 HALO-BAK molecules per assembly (Figure S2C-G). We correlated the stoichiometry of individual HALO-BAK assemblies with their shape (classified as line, arc and ring structures) by correlating the data from photon-counting confocal and STED microscopy (Figure 2A-B). Contrary to our expectation that the HALO-BAK assembly shape would depend on oligomerization state, we did not find significant differences in BAK stoichiometry between line, arc and ring structures (Figure 2B). Despite the broad distribution, the median of BAK stoichiometry was about 100 HALO-BAK molecules in all three categories (Figure 2B and D-F). This suggests that the oligomeric state of BAK does not define the spatial arrangement of its assemblies, implying that it is likely determined by another mechanism.

**Figure 2:**
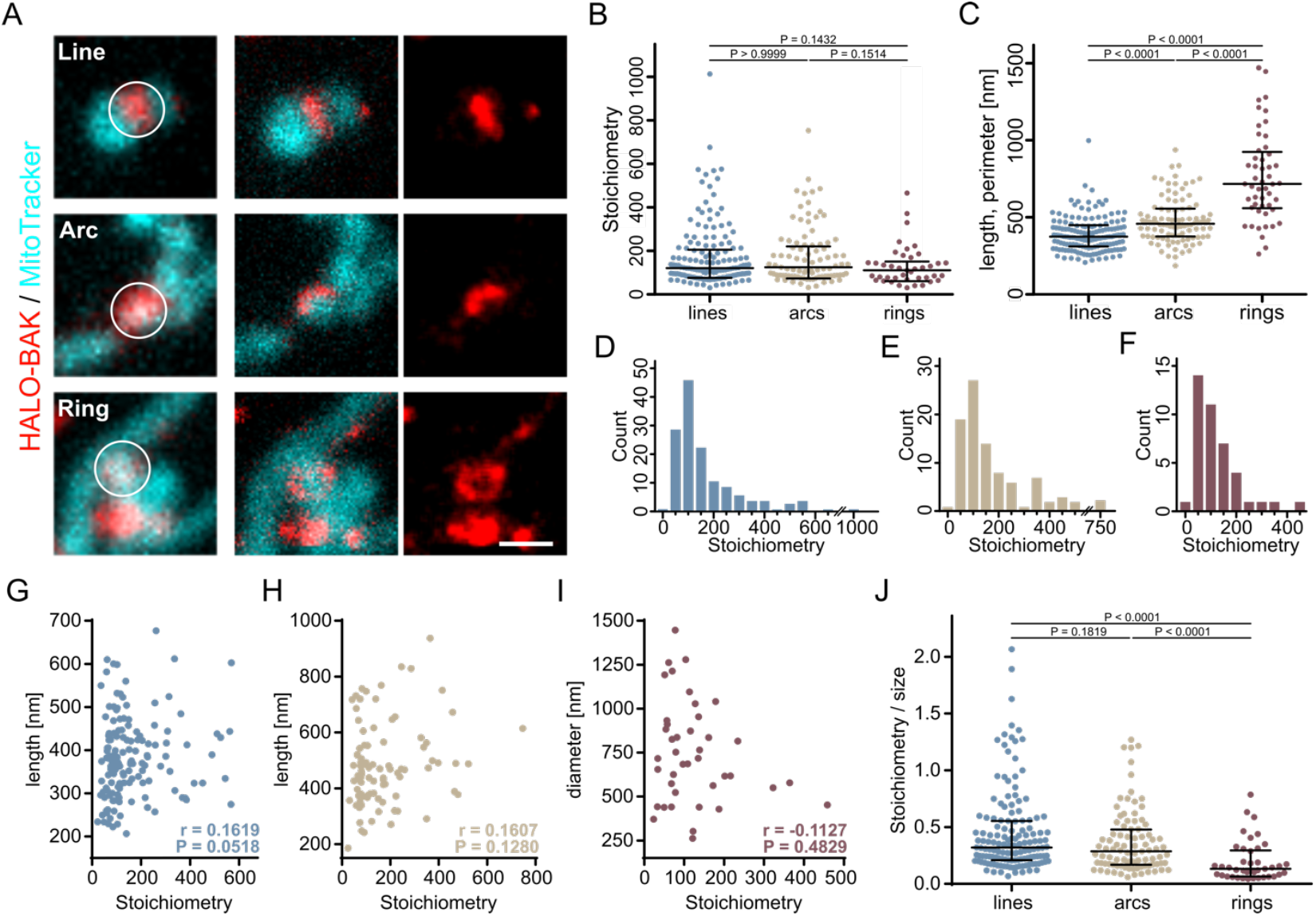
Oligomerization does not determine the supramolecular arrangement of individual BAK assemblies. A) Gallery of representative line, arc and ring structures of BAK (red) visualized using photon-counting confocal (left panel) and STED microscopy (middle and right panel). Mitochondria were visualized using MitoTracker orange (cyan, confocal). Circles indicate regions of interest for stoichiometry quantification. Scale bar 500 nm. B, C) Quantification of the stoichiometry of line, arc, and ring structures (B) as well as the length of line and arc and the perimeter of ring structures (C) formed by BAK. Values correspond to individual structures (individual data points) as well as the median (line) ± interquartile range (whiskers) of n = 145 lines, n = 91 arcs, and n = 41 rings for stoichiometry (B) and n = 154 lines, n = 98 arcs, and n = 50 rings for size measurements (C). Significance levels were determined by non-parametric one-way ANOVA (Kruskal-Wallis test) and Dunn’s multiple comparison test. P indicates multiplicity adjusted P values (family-wise significance and confidence level set to 0.05). D-F) Frequency distribution of the stoichiometry of line (D), arc (E), and ring (F) structures. G-I) Correlation between stoichiometry and the length of lines (blue, G), arcs (beige, H) and ring perimeter (burgundy, I). Values correspond to individual structures (individual data points) of n = 145 lines, n = 91 arcs, and n = 41 rings. r indicates the Spearman correlation coefficient, P indicates the two-tailed P value for each individual category. J) Stoichiometry/size-ratio of individual assemblies sown in (G-I). Values are shown for individual BAK assemblies (individual data points) and the median (line) ± interquartile range (whiskers) of n = 151 lines, n = 91 arcs, and n = 41 rings. Significance levels were determined by non-parametric one-way ANOVA (Kruskal-Wallis test) and Dunn’s multiple comparison test. P indicates multiplicity adjusted P values (family-wise significance and confidence level set at 0.05). All data presented are representative of n = 8 independent experiments.

HALO-BAK structures varied widely in size, ranging from less than 200 nm to over 1.4 µm in length/perimeter, with significant differences between them. With a median length of approximately 400 ± 130 nm, lines were smaller than arcs, which have a median length of approximately 500 ± 200 nm. Rings were larger than both lines and arcs, with a median perimeter of approximately 700 ± 400 nm, corresponding to a median ring diameter of approximately 200 ± 100 nm (Figure 2C). The size distributions align well with previous studies ^9,12^. Yet, we found no significant correlation between the number of HALO-BAK molecules and the size of the structures (Figure 2G-J). The lack of correlation between stoichiometry and size of individual BAK assemblies found here indicates that BAK stoichiometry does not (solely) determine the supramolecular structural organization and size of BAK apoptotic pores (we cannot discard a role for BAX co-oligomerization, not quantified in our experiments).

### HALO-BAK assemblies arrange along MOM openings at apoptotic pores

Given the toroidal nature of the apoptotic pore, we reasoned that other factors such as the physical properties of the membrane environment might contribute to define the pore size and shape. To explore this possibility, we correlated the shape of the supramolecular assemblies detected by STED microscopy with their mitochondrial environment visualized by ET. Generally, we found a strong correlation between the shape of the HALO-BAK nanoassembly with the position and type of MOM opening (Figure 3). We categorized MOM discontinuities (pores) in the EM images according to their appearance and location at mitochondria as i) discontinuities at the tips, ii) openings with MIM extrusion, iii) discontinuities along the lateral axes, or iv) at putative post-fission sites, where two mitochondrial fragments (and ER membranes) were in close proximity (Figure 3). MOM discontinuities appeared most frequently at mitochondrial tips (n = 39, Figures 3A and 5C) and as openings with MIM extrusion (n = 17, Figures 3B and 5D). Lateral discontinuities (n = 6, Figures 3C and 5E) and at putative post-fission sites (n = 2, Figures 3D and 5F) were comparably rare. These findings suggest that mitochondrial tips may be more favorable locations for apoptotic pore formation.

**Figure 3:**
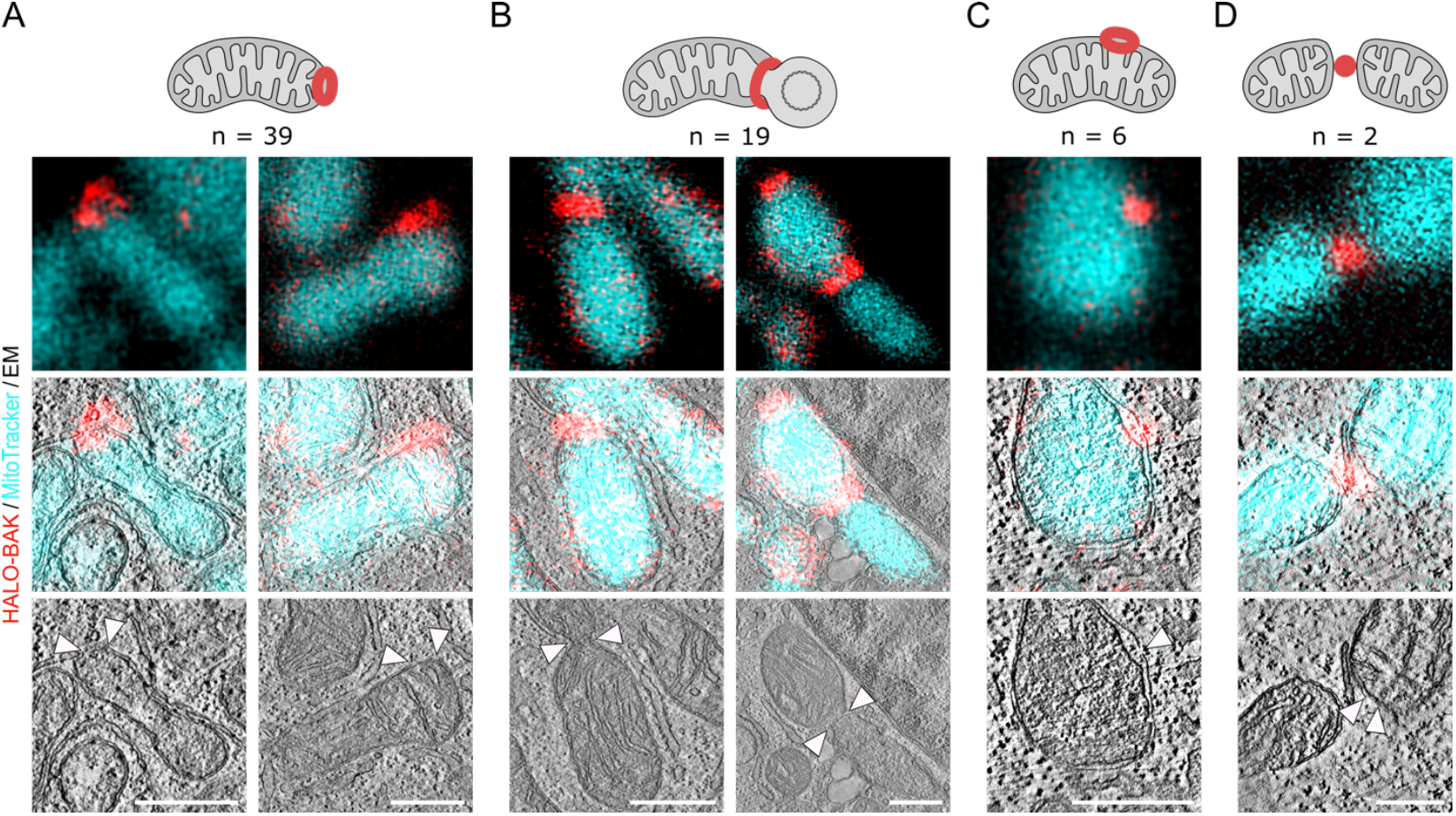
Organellar distribution of BAK pores at the MOM. A) MOM opening at the tip of the mitochondrion, B) MOM opening with extrusion of the MIM, C) MOM opening at the side of the mitochondrion, and D) MOM opening at a potential post-fission site, as indicated by the schematic representation (BAK shown in red). Microscopy images show representative STED microscopy (top), STED-CLEM overlay (middle) and TEM (bottom) examples of mitochondria from U2OS HALO-BAK cells one hour after apoptosis induction labeled with JFX650 Halo-Tag ligand (red, STED) for each category. Mitochondria were labeled with MitoTracker Orange (cyan, confocal). Arrowheads indicate the position of the MOM discontinuity. n indicates the number of cases assigned to the corresponding category out of a total of n = 6 independent experiments. Scale bar 500 nm.

### Biophysical forces promote MIM extrusion and apoptotic pore growth

CLOSE microscopy allowed us to document cases of MIM permeabilization and extrusion through MOM openings and to associate them apoptotic BAK assemblies of different shapes, although with limited statistics. To quantify the frequency MIM extrusion, we implemented an assay based on the visualization of the MOM and the mitochondrial matrix together with HALO-BAK in apoptotic cells under caspase inhibition fixed 1h after treatment, using confocal and STED microscopy (Figure 4A-B). Despite approximately 40% of mitochondria with microscopically intact MOM, MOM openings were associated with MIM extrusion in 39% of mitochondria, whereas only 21% of mitochondria showed MOM openings without MIM extrusion (Figure 4C-D), indicating the widespread nature of this event. Of note, we found BAK localized to one or both edges of the MOM opening but not to the MIM (Figure 4A-B).

**Figure 4:**
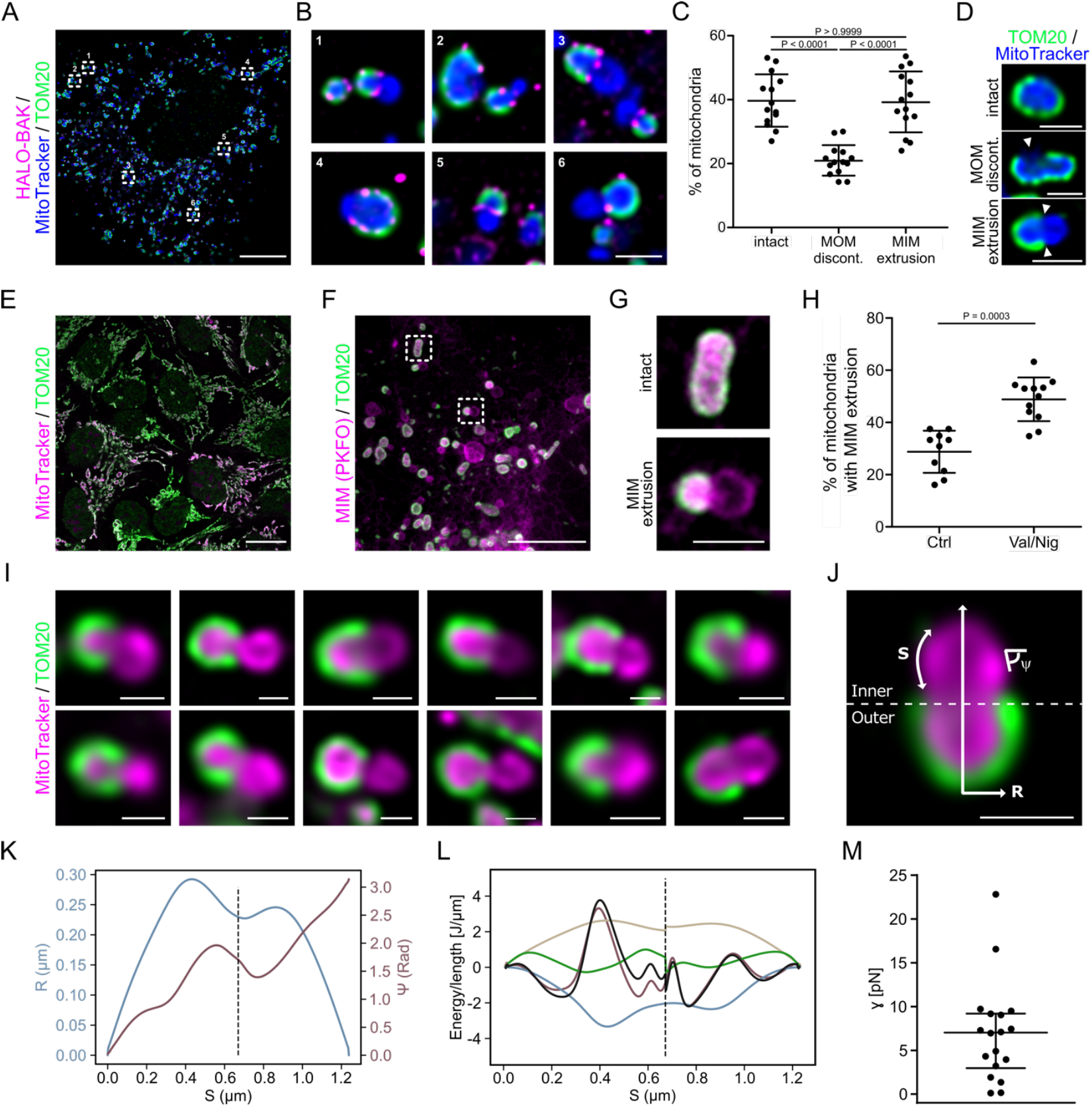
Mechanical forces promote MIM extrusion through BAK pores in the MOM. A) Representative deconvolved confocal/STED microscopy image of U2OS HALO-BAK cells labeled with JFX650 HALO-Tag ligand (magenta, STED) one hour after apoptosis induction. The MOM was stained by immunolabeling against TOM20 (green, confocal) and the mitochondrial matrix was visualized by MitoTracker Orange (blue, confocal). Numbered rectangles indicate cropped regions in (B). Scale bar 10 µm. B) Enlarged images of cropped regions of the overview image shown in (A) as indicated. Scale bar 1 µm. C) Quantification of apoptotic mitochondria with intact MOM (intact) versus mitochondria with discontinuity of the MOM (MOM discont.) or mitochondria with extrusion of MIM through MOM openings (MIM extrusion). Values correspond to the percentage of mitochondria of individual cells (individual data points) as well as the mean (line) ± SD (whiskers) of n = 14 cells. Significance levels were determined by non-parametric one-way ANOVA (Kruskal-Wallis test) and Dunn’s multiple comparison test. P indicates multiplicity adjusted P values (family-wise significance and confidence level set to 0.05). D) Representative confocal microscopy images of the categories quantified in (C) of U2OS HALO-BAK cells prepared as described in (A). Scale bar 1 µm. E) Representative confocal microscopy overview image of valinomycin/nigericin-induced osmotic mitochondrial matrix swelling in healthy U2OS HALO-BAK cells. The MIM/mitochondrial matrix was visualized by MitoTracker Orange (magenta) and the MOM was stained by immunolabeling against TOM20 (green). Scale bar 20 µm. F) Representative confocal/STED microscopy image of U2OS HALO-BAK cells transiently transfected with TOM20-SNAP labeled with JFX650 SNAP-tag ligand (green, deconvolved confocal) one hour after apoptosis induction. The MIM was labeled with PKFO (magenta, STED). Dotted rectangles indicate cropped regions in (G). Scale bar 5 µm. G) Enlarged images of cropped regions of the image shown in (F) as indicated, showing exemplary mitochondria with intact MOM (intact) or mitochondria with MIM extrusion through MOM openings (MIM extrusion). Scale bar 1 µm. H) Quantification of mitochondria with MIM extrusion through MOM openings in apoptotic U2OS HALO-BAK control (Ctrl) cells or apoptotic U2OS HALO-BAK cells treated with valinomycin/nigericin (Val/Nig) to induce osmotic swelling of the mitochondrial matrix. Values correspond to the percentage of mitochondria of individual cells (individual data points) as well as the mean (line) ± SD (whiskers) of n = 10 (ctrl) and n = 12 (Val/Nig) cells. Significance levels were determined by unpaired nonparametric Student’s t-test (Mann-Whitney test, P value as indicated). Data are representative of n = 5 (panels A-D) and n = 4 (panels F-H) independent experiments. I) Gallery of hourglass-shaped MIM extrusion examples from U2OS HALO-BAK cells one hour after apoptosis induction. The MOM was stained by immunolabeling against TOM20 (green, deconcolved) and the mitochondrial matrix was visualized by MitoTracker Orange (magenta). Scale bar 500 nm. J) Representative axisymmetric image of a MIM extrusion as described in (I). The mitochondria are aligned so that the longitudinal axis is orthogonal to the axis of the opening pore separating the regions of exposed MIM (inner) and MIM contained in MOM (outer, dashed line). The parametrization of the local radius R and the tangent angle ψ as a function of the arc length S is presented. Scale bar 500 nm. K) Local mitochondrial radius R and tangent angle ψ as a function of the arc length S, in blue and burgundy, respectively. L) Minimized energy of the experimental mitochondria presented in (J). In black, the sum of the energies is plotted as a function of the arc length S. The lateral tension, pressure, bending, and line energies are plotted as a function of S, in beige, blue, burgundy, and green, respectively. M) Pore rim line tension γ obtained from the minimization routine of n = 18 mitochondria from n = 9 individual cells. Data are presented as the line tension values for individual mitochondria (individual data points) as well as the median (line) ± interquartile range (whiskers). Data are representative of n = 3 independent experiments.

Given the lack of correlation between oligomerization and pore size, we hypothesized that MIM extrusion through apoptotic MOM pores, promoted by MIMP-induced matrix swelling, might mechanically contribute to pore expansion. To test this hypothesis, we first evaluated whether matrix swelling promotes MIM extrusion. To this aim, we sought to induce osmotic swelling in the mitochondrial matrix by treatment with the ionophores valinomycin and nigericin (Val/Nig) ^35^, which we confirmed by confocal microscopy in healthy cells (Figure 4E). When applied to apoptotic cells, we found a significantly higher ratio of mitochondria with MIM extrusion in Val/Nig-treated cells compared to control untreated apoptotic cells (Figure 4F-H). These results indicate that increased mitochondrial matrix swelling (due to osmotic pressure) promotes MIM extrusion, uncovering a role of biophysical forces in the mechanism of MIM extrusion.

Interestingly, we observed that MIM extrusions through restricted openings of the MOM were squeezed at the site of the MOM opening adopting an hourglass shape (Figure 4B, D, G and I). This membrane deformation from the spherical shape of minimal energy reveals that the MOM opening exerts a force on the extruding MIM resulting from a line tension at the MOM edge that tends to close the pore. Such membrane arrangements also show that the MIM in turn counterbalances this force and keeps the MOM pore open.

By using an adapted version of the quantitative “neck model” geometrical analysis proposed by Baumgart et al. in ^36^, we estimated the line tension at the edge of the apoptotic pore directly in mitochondria of apoptotic cells. The analysis is based on the minimization of the energy contributions associated with membrane curvatures and tensions, transmembrane pressure, and line tension consistent with the measured membrane geometry based on the conservation of a zero value Hamiltonian ^36,37^. For our analysis, we used images of axisymmetric hourglass-shaped MIM extrusions through MOM pores and aligned the mitochondrial longitudinal axis orthogonal to the open MOM pore axis. By tracing the MIM outline in the image, the local radius R and the local tangent angle ψ can be obtained as a function of the distance of the arc length S of the traced MIM (Figure 4J-K). Using a condition in which we minimize the energy of the Hamiltonian (by modulating the values of the membrane bending modulus, lateral tension and outer excess pressure, see Methods section) together with the experimental data obtained from the images, we estimated a line tension of *γ* = 7 ± 6 pN at the apoptotic pore edge on the MOM (Figure 4L-M). This is in good agreement with the line tension estimated for BAX α-helix 5-mediated pores in model membranes ^18^.

### Role of MIM confinement and permeabilization in extrusion through the MOM

We performed membrane dynamics simulations to gain further insights into the MIM extrusion process and to quantify the associated forces. We modeled the MIM as a triangulated fluid-elastic membrane with *cristae*-like structures densely packed into a pill-shaped container mimicking the MOM. To mimic MOMP, we opened a circular hole of fixed size at one end of the MOM container through which the MIM could escape.

In our simulations, we observed spontaneous MIM extrusion for apoptotic pores at the MOM whose radius exceeded 50% of the mitochondrial diameter (Figure 5A and S4A; Supplementary Movie 2). This evagination was associated with *cristae* rounding, which appeared to push the MIM through the MOM opening. Over time, we also observed *cristae* opening or merging with each other. Driving the MIM remodeling and extrusion in the simulations was a substantial reduction in the membrane bending energy.

**Figure 5:**
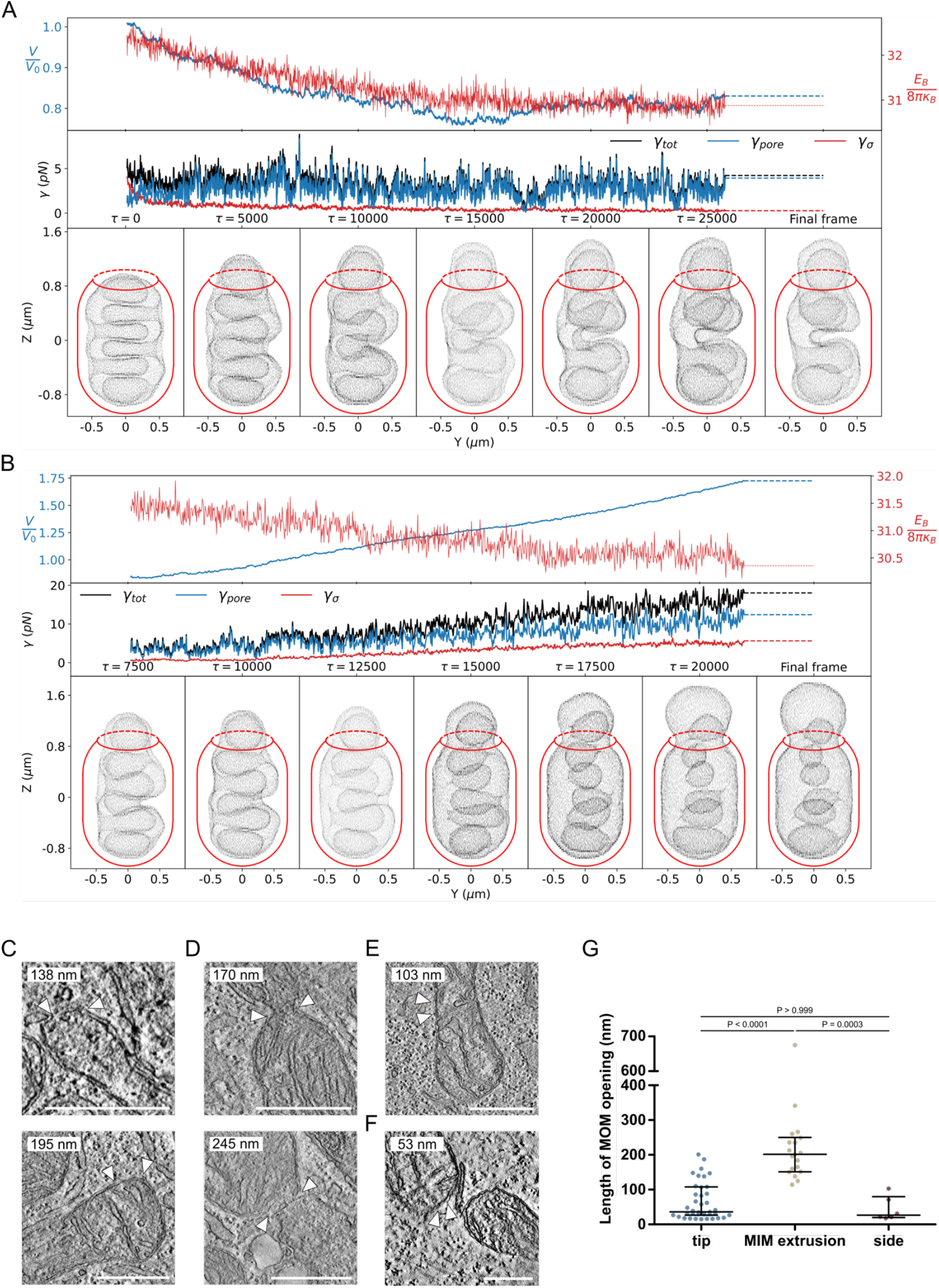
Interplay between MIM extrusion and the apoptotic pore. A-B) Membrane dynamic simulations of MIM extrusion through the apoptotic pore at the MOM. MIM (grey in bottom panel) contained in MOM (red schematic outline) with a pore of size *α* = 0.75 corresponding to a relative diameter of 0.682 and a bending rigidity of κ_B_ = 10 *k*_B_*T* without (A) and with osmotic inflation after time *τ* = 7500 (B). The bottom panels show snapshots at the indicated time points, the top panels the membrane bending energy *E*_B_ (red) and the mitochondrial matrix volume relative to that of the starting structure (blue). The center panel shows the effective line tension created by the evaginating membrane in the apoptotic pore (see Methods for components). C-G) Quantification of the size of individual MOM openings. C-F) Exemplary EM images of single mitochondria with MOM discontinuity at the tip of the mitochondrion (C), MOM opening with MIM extrusion (D), MOM discontinuity at the side of the mitochondrion (E), and at a potential post-fission site (F). Arrowheads indicate the position of the MOM discontinuity and the measurement position. The size of the measured opening is indicated in each image. Scale bar 500 nm. G) Quantification of the size of individual MOM openings according to the categories described in (C-E). Values correspond to individual measurements (individual data points) as well as the median (line) ± interquartile range (whiskers) of n = 39 MOM openings at the tip, n = 19 MOM openings with MIM extrusion, and n = 6 MOM openings at the side of the mitochondrion from n = 6 independent experiments. Significance levels were determined by non-parametric one-way ANOVA (Kruskal-Wallis test) and Dunn’s multiple comparison test. P indicates multiplicity adjusted P values (family-wise significance and confidence level set to 0.05).

As the MIM extrudes, it exerts an expansive force on the rim of the apoptotic pore, which we quantified in terms of an effective line tension (see Methods). For the pore in Figure 5A, with a relative diameter of *≈* 70% of the mitochondrial diameter and a low membrane bending rigidity of κ_*B*_ = 10 *k*_*B*_*T* ^38^, we obtained line tensions in the range of *γ*_tot_ *≈* 3 − 5 pN. For stiffer membranes, the line tensions were somewhat higher (*γ*_tot_ *≈* 10 − 15 pN for κ_B_ = 60 kT; Figure S4B). For larger pores, the effective line tension was reduced (Figure S4C). These line tensions are in quantitative agreement with the tensions of *γ ≈* 7 ± 4 pN deduced from the membrane shapes in the fluorescence microscopy images.

In the initial phase of MIM extrusion, we found that the matrix volume tended to decrease, associated with *cristae* rounding. However, as MIM extrusion progressed, we also observed volume increases (Figure 5A top panel). More dramatic volume increases were observed for the stiffer membranes with the larger pore, of diameter comparable to the mitochondrial diameter, as MIM extruded more and *cristae* opened up outside the MOM (Figure S4D).

To mimic the conditions of MIMP in apoptosis ^24,25^, we performed simulations with matrix osmotic swelling as a result of MIM leaks. In line with the microscopy data, we found that matrix swelling accelerated MIM extrusion (Figure 5B; Supplementary Movie 3), which also resulted in larger effective line tensions on the apoptotic pore at the MOM, reaching values *>* 10 pN. Interestingly, even substantial osmotic forces did not result in significant MIM extrusion through a small (*<* 50% relative diameter) and static MOM pore (Figure S4E). However, the observed buildup in line tension would likely expand the apoptotic pore over time.

According to these results, we would expect that MIM extrusion events would be associated with larger apoptotic pores. To test this, we determined the length of individual MOM discontinuities measured as the maximum linear distance of interruption between the two ends of the MOM in the EM tomogram (Figure 5C-G). While individual examples of MOM discontinuities varied widely in length (Figure 5C-F), MOM openings associated with MIM extrusion were significantly larger than MOM discontinuities at the mitochondrial tip or side (Figure 5G), in agreement with the simulation predictions. These results indicate that MIM extrusion promotes apoptotic pore growth, impacting its structural organization.

## Discussion

Here we present CLOSE microscopy, a multi-correlative approach that allows dissecting the oligomeric state of individual, discrete protein membrane complexes, based on photon-counting confocal imaging, and their supramolecular structural arrangement, imaged with STED microscopy, in relation to their cellular environment, visualized by electron tomography. It provides an easily implementable pipeline that brings together these three types of structural information and allows the study of focal protein complexes with unprecedented detail, opening new avenues for uncovering their molecular mechanisms. In future developments, it can be extended to live-cell super-resolution microscopy and combined with cryo-EM.

We apply CLOSE microscopy to study the assembly of the apoptotic pore and its relation with mitochondrial alterations in apoptotic cells (Figure 1). Our findings indicate that apoptotic pore size and shape are not determined, at least solely, by BAK oligomerization, although we cannot disentangle here the role of BAX on this process. We find that HALO-BAK oligomers delineate the apoptotic pore openings at the MOM, suggesting that it is the membrane opening that defines the spatial arrangement of BAK assemblies, which would also explain the heterogeneity in BAK nanoassemblies. The majority of MOM pores were located at tips, suggesting a preferential site for apoptotic pore assembly. Since dynamin-related protein 1 (DRP1) is functionally linked to apoptotic pore formation ^39–42^, it is reasonable to assume that the BAK assemblies and MOM pores at mitochondrial tips may correspond to post-fission sites, as proposed in ^43^. Remarkably, we could visualize a few examples of pores at the MIM still contained within the MOM, which raises the possibility of a kinetically regulated mechanism governing MIMP either within the mitochondrion or following MIM extrusion.

The finding that induced osmotic mitochondrial matrix swelling promoted MIM extrusion indicates that biophysical forces contribute to the MIM extrusion process. In addition, the hourglass shape of the MIM extruding through the MOM pores revealed constriction by the MOM resulting from line tension at the MOM opening edge. This implies that the force exerted by unfolding and extrusion of the MIM through MOM openings contributes to pore widening independent of BAK oligomerization, which is reasonable considering the lipid/protein nature of the apoptotic pore. Accordingly, we could qualitatively and quantitatively recapitulate MIM extrusion from mitochondria containing apoptotic pores at the MOM in membrane dynamic simulations. Above a critical pore size, the MIM readily evaginated from the MOM in a process energetically driven by a reduction in bending energy of the MIM, which is densely packed into *cristae* inside the MOM in healthy cells. Remarkably, the effective line tensions on the apoptotic pore calculated directly from the mechanical forces in the membrane dynamic simulations agree well with the line tensions deduced from the membrane shapes seen by microscopy. In line with the wet lab experiments, osmotic swelling of the matrix further accelerated the matrix extrusion in the membrane dynamic simulations. It also induced a larger line tension at the apoptotic pore. While the rim was static in our simulations, in apoptotic mitochondria the pore is dynamic, so that the increased line tensions seen under osmotic swelling will likely further expand the pore unless balanced by contracting forces, in line with the experimental data. Any pore expansion would thus result in a positive feedback loop, as larger pores would substantially speed up MIM extrusion.

Collectively, our findings reveal that unfolding, swelling, and extrusion of the MIM mechanically stabilize the apoptotic pore and promote its growth by counteracting the line tension at the edge of the pore. They thus support a mechanism in which oligomerization of BAK (and BAX) at the pore rim would initially open the pore and drive its expansion. Apoptotic pore growth would then enable the extrusion of the MIM, further promoted by matrix swelling following MIMP. Beyond this point, MIM extrusion would enhance apoptotic pore expansion at the MOM by counteracting the line tension at the pore rim. If this tension is at least partially balanced by the pressure of the extruding MIM, additional recruitment of BAK (and BAX) to the pore rim might shield exposed hydrophobic membrane regions and further stabilize the pore. This model could explain the lack of correlation between BAK stoichiometry and pore size, as well as why often large pores are not fully lined by BAK (and BAX) molecules.

Functionally, this may have consequences for determining the irreversibility of MOMP as well as for MIMP and the downstream inflammatory signaling triggered by mitochondrial contents release. On the one hand, while antiapoptotic BCL2 family members are capable of disassembling BAX oligomers to inhibit pore formation ^44^, MIM extrusion through the apoptotic pore involves reorganization of mitochondrial membranes to an extent that MOMP becomes energetically irreversible. This would likely leave the elimination of the permeabilized mitochondrion, for example by mitophagy ^45^, as the only mechanism to limit the cellular effects of mitochondrial permeabilization, thereby contributing to the transition from sublethal to lethal MOMP. On the other hand, exposure of the MIM to cytosolic components, both through the apoptotic pore and upon extrusion, would make it accessible to potential mediators of MIMP. The extruded MIM is no longer protected by the MOM, increasing its fragility and making it susceptible to rupture by mechanical stress, as an alternative mechanism of MIMP. Our findings uncover a previously unrecognized interplay between the MOM and the MIM that governs the dynamics of the apoptotic pore through biophysical forces, which may have consequences for MOMP irreversibility and downstream inflammatory signaling.

## Materials and Methods

### Cell culture

U2OS wild type, U2OS HALO-BAK, U2OS NUP96-HALO and HEK 293T cells were cultivated at 37 °C and 5% CO_2_ in DMEM supplemented with 10% FBS and 1% penicillin/streptomycin (Invitrogen, Germany). The U2OS NUP96-HALO (U-2_OS-CRISPR-NUP96-Halo_clone_no252) CRISPR cell line was obtained from CLS Cell Lines Service GmbH, Eppelheim, Germany. All cell lines were tested mycoplasma negative.

### Generation of U2OS HALO-BAK knock-in cell line using CRISPR/Cas9

CRISPR/Cas9-mediated knock-in of HALO-Tag into the *BAK* locus directly upstream of the start codon was performed by a combinatorial approach using lentiviral transduction of the lentiCRISPR v2 (Addgene #52961, ^46^) vector containing the SpCas9 enzyme and a single guide RNA (sgRNA) targeting *BAK* and transfection of a homology-directed repair (HDR) template containing 800 bp *BAK* homology arms and the HALO-Tag expression cassette into wild-type U2OS cells (Figure S1A). Genome-edited single cell clones were isolated and validated by sequencing of the *BAK* locus, immunoblotting, and fluorescence microscopy (Figure S1B-F).

### sgRNA design and cloning of lentiCRISPR v2 vector

sgRNAs were designed on the human genomic *BAK* locus (*BAK1*, ENSG00000030110) near the start of the *BAK* open reading frame using the CRISPR guide RNA design tools Benchling and CRISPOR (Table 1). Suitable 20 bp sgRNAs with a NGG PAM motif were filtered based on the predicted on-target ^47^ and off-target ^48^ scores compared to the human reference genome (GRCh38 (hg38, Homo sapiens), Genome Reference Consortium) considering the described genome variants (dbSNP148, ^49^). The selected sgRNA (sgRNA6: 5’-TGGAGGACGGGATCAGCCTG-3’) was cloned into the lentiCRISPR v2 lentiviral transfer vector using a corresponding synthetic DNA oligonucleotide set (sgRNA6-forward: 5’-caccgTGGAGGACGGGATCAGCCTG-3’ and sgRNA6-reverse: 5’-aaacCAGGCTGATCCCGTCCTCCAc-3’) according to an adapted version of ^50^.

**Table 1:**
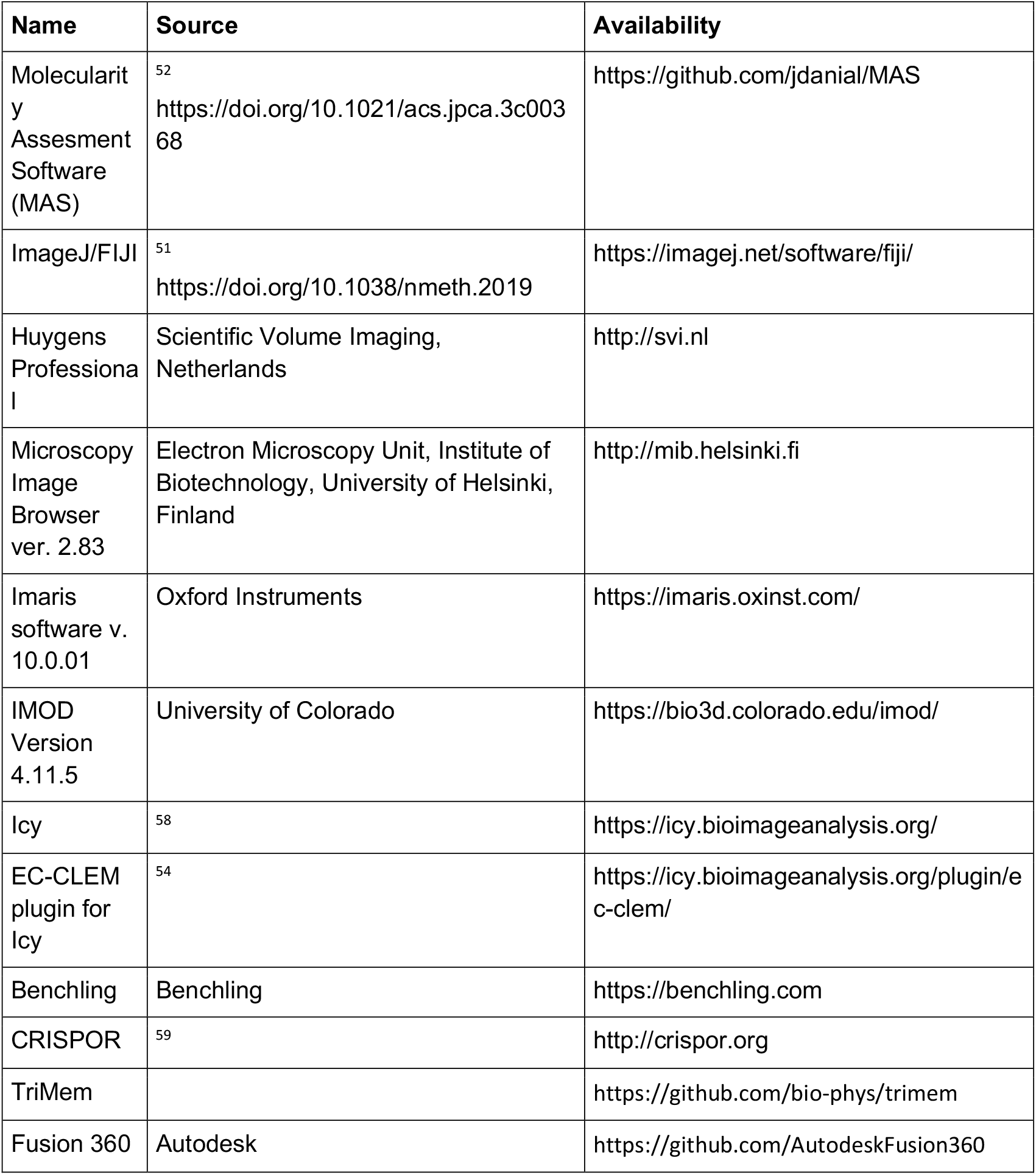
Software.

| **Name** | **Source** | **Availability** |
| --- | --- | --- |
| Molecularity Assessment Software (MAS) | <sup>52</sup><br>https://doi.org/10.1021/acs.jpca.3c00368 | https://github.com/jdanial/MAS |
| ImageJ/FIJI | <sup>51</sup><br>https://doi.org/10.1038/nmeth.2019 | https://imagej.net/software/fiji/ |
| Huygens Professional | Scientific Volume Imaging, Netherlands | http://svi.nl |
| Microscopy Image Browser ver. 2.83 | Electron Microscopy Unit, Institute of Biotechnology, University of Helsinki, Finland | http://mib.helsinki.fi |
| Imaris software v. 10.0.01 | Oxford Instruments | https://imaris.oxinst.com/ |
| IMOD Version 4.11.5 | University of Colorado | https://bio3d.colorado.edu/imod/ |
| Icy | <sup>58</sup> | https://icy.bioimageanalysis.org/ |
| EC-CLEM plugin for Icy | <sup>54</sup> | https://icy.bioimageanalysis.org/plugin/ec-clem/ |
| Benchling | Benchling | https://benchling.com |
| CRISPOR | <sup>59</sup> | http://crispor.org |
| TriMem |  | https://github.com/bio-phys/trimem |
| Fusion 360 | Autodesk | https://github.com/AutodeskFusion360 |

### Design and generation of the HDR template

Homology regions to the *BAK* locus (800 bp upstream and downstream of the start codon, omitting the start codon) were amplified from genomic DNA isolated from wild-type U2OS cells and cloned into pUC19 (Addgene #50005) including a multiple cloning site between the homology regions using Gibson assembly. The HALO-Tag sequence (including the start codon and a (GGS)_3_ linker) was amplified from pSems-HALO7-tag and cloned in between the homology regions into the pUC19-BAK-HDR vector by restriction cloning.

### Virus production

To generate transgenic lentiviruses, HEK 293T cells were seeded in a 6-well plate at a density of 8 × 10^5^ cells per well one day prior to transfection. The cells were transfected with 1.5 µg of the lentiCRISPR v2-BAK-gRNA6 lentiviral transfer vector, 650 ng of the psPAX2 lentiviral packaging vector (Addgene #12260), and 350 ng of the pMD2.G lentiviral envelope vector (Addgene #12259) using Lipofectamine™ 2000 (ThermoFisher). The cells were cultured for 4 days to allow virus production. On day 4, the virus-containing supernatant was collected, centrifuged at 1250 rpm for 5 minutes, and filtered through a 0.45 µm filter. Polybrene at a final concentration of 8 µg/ml was added to the viral supernatant prior to infection to increase transduction efficiency.

### Cellular delivery and single cell sorting

Wild-type U2OS cells were transduced with the lentivirus containing the SpCas9-BAK-gRNA6 transfer cassette and co-transfected with 2 µg of the pUC19-BAK-HALO-HDR construct using Lipofectamine™ 2000. One week after transfection/transduction, the cells were stained with 300 nM Janelia Fluor X 650 HaloTag® Ligand (Janelia Materials) for 45 minutes at 37°C. After staining, the cells were washed with DMEM and PBS to remove unbound dye. The cells were then analyzed by flow cytometry and individual double-positive cells (positive for both JFX650 and GFP signal) were isolated. Gating was performed to exclude dead cells using the DAPI channel to ensure that only viable cells were collected. Single-cell clones were expanded and validated by PCR genotyping of genomic DNA, immunoblotting, and fluorescence imaging.

### Validation by Western blot

Cells were harvested using trypsin-EDTA (Sigma), resuspended in culture medium, and collected by centrifugation at 300 × g for 5 minutes at 4 °C. The cell pellets were washed twice with ice-cold PBS. For lysis, the pellets were resuspended in RIPA lysis buffer (50 mM Tris/HCl, pH 8.0, 150 mM NaCl, 1% (v/v) Triton™ X-100, 0.5% (w/v) sodium deoxycholate, 0.1% (w/v) SDS), incubated on ice for 20 minutes, and centrifuged at 20,000 × g for 20 minutes at 4 °C to remove cellular debris. Protein concentration was determined using the Bradford protein assay (Bio-Rad) according to the manufacturer’s protocol. A total of 50–100 µg of protein was boiled in SDS-PAGE sample buffer (62.5 mM Tris/HCl, pH 6.8, 2% (w/v) SDS, 10% (v/v) glycerol, 0.005% (v/v) β-mercaptoethanol, 0.01% (w/v) bromophenol blue) for 5 minutes at 95°C prior to SDS-PAGE. Proteins were transferred to a nitrocellulose membrane (Trans-Blot Turbo, Bio-Rad), and equal sample loading was confirmed using Ponceau S staining. The blots were washed with TBST (50 mM Tris/HCl, pH 7.5, 150 mM NaCl, 0.1% (v/v) Tween 20) and blocked with 5% (w/v) low-fat milk in TBST for 60 minutes. The membranes were incubated with a 1:1000 (v/v) dilution of rabbit α-BAK primary antibody (D4E4, CST, #12105) in 5% (w/v) milk in TBST at 4 °C for 16 hours. After three washes with TBST (5 minutes each), the membranes were incubated with a 1:10,000 (v/v) dilution of α-rabbit IgG-HRP secondary antibody (JIR, #111-035-003) in 5% (w/v) milk in TBST for 1 hour at room temperature. Blots were washed three times with TBST and developed using SuperSignal™ West Pico PLUS chemiluminescent substrate (Thermo Scientific), followed by detection on the Fusion SL Gel Chemiluminescence Documentation System (Vilber Lourmat). Images were adjusted for brightness and contrast, and cropped using Fiji/ImageJ (Table 1, ^51^).

### Validation by confocal microscopy and quantification of cell death kinetics

U2OS HALO-BAK cells were seeded in a glass-bottom 8-well μ-slide (IBIDI) at a density of 2×10^4^ cells per well one day prior to the experiment. Cells were stained with 300 nM Janelia Fluor X 650 HaloTag® Ligand for 1 hour at 37 °C to label BAK and with 150 nM MitoTracker™ Green FM (ThermoFisher, #M7514) to visualize mitochondria or with 150 µM tetramethylrhodamine ethyl ester (TMRE, ThermoFisher) for 20 minutes at 37 °C to measure mitochondrial depolarization. After labeling, the cells were washed twice with DMEM and incubated for 30 - 50 minutes at 37 °C to remove unbound dye. The localization of BAK in healthy conditions and the formation of BAK apoptotic foci and mitochondrial depolarization (loss in TMRE signal) during apoptosis were measured using an inverted laser scanning STED microscope (TCS SP8 STED 3x, Leica, Wetzlar) equipped with a PL Apo 100x/1.40 Oil STED_Ornage_ objective (Leica, Wetzlar) and a tunable white light laser source (Leica, Wetzlar). MitoTracker™ Green FM was excited at a wavelength of 488 nm, JFX650 at 640 nm, and TMRE at a wavelength of 561 nm. The fluorescence emission signal of JFX60 and MitoTracker™ Green FM of JFX650 and TMRE was detected line sequentially on GaAsP detectors. Single confocal snapshots of individual cells were acquired to validate BAK expression and localization. To measure cell death kinetics, confocal time-lapse tile scans were acquired with a time resolution of 1 minute for a duration of 60 minutes after apoptosis induction using 10 µM ABT737 (Hölzel, #HY-50907), 10 µM S63845 (Hölzel, #HY-100741), and 10 µM Q-VD-OPh (Hölzel, #HY-12305g) in phenol red free DMEM at 37 °C and 5% CO_2_. The average TMRE emission signal intensity was measured over time in individual cells using InageJ/Fiji (Table 1, ^51^) and fitted with a one-phase exponential decay function to calculate the time point of 50% TMRE signal loss (t_50_). The population distribution of the individual t_50_ values was fitted with a Gaussian function to derive the mean lag-time of 50% TMRE signal loss after apoptosis induction. For the further stoichiometry quantification, a time point 20-30 minutes after 50% TMRE signal loss (60 minutes after cell death induction) was chosen to ensure that the majority of cells underwent MOMP.

### CLOSE microscopy

#### Sample preparation for light microscopy

U2OS HALO-BAK and U2OS NUP96-HALO cells were seeded in a glass-bottom imaging dish (MatTek, #P356-1.5-14-C) coated with a carbon finder grid using a mask (Leica, #16770162) and an ACE 200 carbon coater (Leica) at a density of 10^5^ cells per dish one day before the experiment. U2OS NUP96-HALO cells are required as a calibration sample for ratiometric stoichiometry quantification and were treated in the same way as HALO-BAK cells throughout the preparation procedure unless indicated differently. The cells were stained with 300 nM Janelia Flour X 650 (JFX650) HALO-Tag ligand (Janelia Materials) for 1 hour at 37 °C to label BAK (or NUP96) followed by staining with 150 nM MitoTracker™ Orange CMTMRos (ThermoFisher, #M7510) for 30 minutes at 37 °C to visualize mitochondria. The cells were rinsed three times with DMEM and incubated for 1 hour at 37 °C to remove unbound dye. Apoptosis was induced in HALO-BAK cells with 10 µM ABT737 (Hölzel, #HY-50907), 10 µM S63845 (Hölzel, #HY-100741), and 10 µM Q-VD-OPh (Hölzel, #HY-12305g) for 50 – 60 minutes at 37 °C while the cells were monitored repeatedly at the microscope to check the progression of cell death (BAK foci formation). The cells were fixed with 2% glutaraldehyde (Sigma, #G5882) in fixation buffer (100 mM HEPES/KOH pH 7.4 (Carl Roth, #9105.1), 3 mM CaCl_2_ (Sigma, #C7902), 2.5% (w/v) sucrose (Carl Roth, #4621.1), in HPLC water) for 30 minutes at room temperature and 30 minutes at 4 °C. The fixation reaction was quenched with 0.1 M glycine (Carl Roth, #0079.1) in cacodylate buffer (0.1 M sodium cacodylate (Applichem, #A2140.0100) in HPLC water) for 20 minutes at room temperature and the cells were washed three times with 0.1 M cacodylate buffer to remove excess quenching solution. The cells were kept in the dark at room temperature for subsequent microscopy or stored at 4 °C overnight.

#### Photon-counting confocal and STED microscopy

Protein stoichiometry was determined using photon-counting confocal microscopy as described in ^9,31^. Briefly, confocal microscopy images were acquired with an inverted laser scanning STED microscope (TCS SP8 STED 3x, Leica, Wetzlar) equipped with a HC PL APO 100x/1.40 Oil STED_white_ objective (Leica, Wetzlar) and a tunable white light laser source (Leica, Wetzlar). Z-stacks of 8 image planes were collected with an axial spacing of 300 nm, which was determined from point-spread function analysis of fluorescent beads as the maximum spacing required to record at least 80% of the maximum possible intensity from any point source. JFX650 was excited at 640 nm and MitoTracker™Orange CMTMRos at 561 nm. The excitation laser intensity and pixel dwell time were optimized to yield an emission signal detectable in the linear detector range while minimizing photobleaching during the acquisition process. Images were acquired with a pixel resolution of 1024 × 1024 pixel in a region of 46.5 × 46.5 µm (pixel size 45.45 nm) and a pixel dwell time of 2.44 µs (100 Hz scan speed) with 2-fold line averaging. Single emission photons were detected on a hybrid detector in photon counting mode. The NUP96-HALO cells were acquired with the same settings as the HALO-BAK cells for ratiometric stoichiometry analysis as described in ^9,31^. Subsequently, the JFX650 and MitoTracker emission signal was acquired in the HALO-BAK cells in the same (lateral and axial) region of interest in STED (for JFX650, STED de-excitation at 775 nm) and confocal mode (for MitoTracker) with a pixel resolution of 2112 × 2112 pixel (pixel size 22.03 nm) and a pixel dwell time of 588 ns (200 Hz scan speed) with 8-fold line averaging. 2D STED z-stacks were acquired with the same axial spacing as in photon-counting mode. The fluorescence emission signal was detected line sequentially using hybrid detectors with a gating of 0.5 - 6 ns (for the STED image) and 8-fold line averaging. The localization coordinates of the acquired cells on the carbon finder grid were noted for position correlation for electron microscopy image acquisition. Images were adjusted for brightness and contrast and smoothed using ImageJ/Fiji (Table 1, ^51^). Where mentioned, images were deconvolved using Huygens Professional version 19.04 (Scientific Volume Imaging, The Netherlands, Table 1, http://svi.nl).

#### Stoichiometry quantification by ratiometric approach

Single particle detection and brightness analysis was performed on maximum intensity z-projections of photon-counting confocal microscopy images of HALO-BAK using an in-house Python algorithm (now available as an automated Molecularity Assessment Software (MAS, Table 1, ^52^) as described in ^9^. Briefly, single particles were detected in individual 5 × 5 pixel regions of interest (ROIs) based on fluorescence intensity threshold using a Difference of Gaussian detection algorithm. Particle intensities were fitted to a 2D Gaussian function and local background subtraction was performed in a slightly larger ROI around the particle ROI. Importantly, particles with their maximum intensity in the outer z-range were discarded from the analysis, as were particles with overlapping ROIs. To rule out the presence of more than one particle in a ROI, the localized particles were filtered based on the width of the 2D Gaussian. The full width at half maximum (FWHM) of all fitted particles was plotted and ROIs whose FWHM fell outside the 95^th^ percentile of the normal distribution of the entire ROI population were excluded. Using the ratiometric approach, the stoichiometry of a particle of interest S_P_ was determined by ratiometric comparison of its fluorescence intensity I_P_ with the intensity of a reference standard I_S_ of known stoichiometry S_S_ labeled with the same fluorophore as the protein of interest according to the formula S_P_ = I_P_ / (I_S_ / S_S_). The 32-mer HALO-tagged nuclear pore complex protein NUP96, endogenously expressed in U2OS cells ^34^, was used as an internal calibration standard. For each cell, the intensity of the calibration standard I_S_ corresponds to the mean value of the Lognormal-fitted fluorescence intensity distribution individual NUP96-HALO oligomers from at least 5 imaged cells, measured on the same day and with the same parameters as the HALO-BAK samples (Figure S2A-B). For a given cell, the stoichiometry of individual HALO-BAK foci was plotted as a probability density function as well as cumulative counts. Exponential decay fitting was applied to the cumulative distribution to extract the average stoichiometry of HALO-BAK foci per cell (Figure S2C-E).

#### Sample preparation for electron microscopy

After light microscopy, cells were incubated with 1% osmium tetroxide (Science Services, #E19190), 1% potassium ferricyanide (Sigma, #P8131), and 1.25% sucrose (Roth, # 4621.1) in 0.1 M cacodylate buffer (Applichem, # A2140,0100) for 30 minutes at 4 °C. After washing three times with 0.1 M cacodylate buffer for 5 minutes each, samples were dehydrated in an ascending ethanol series (50%, 70%, 90%, and 100% (v/v) ethanol (VWR, #153386F) in ddH2O) for 7 minutes each at 4 °C. Cells were infiltrated with a 50% (v/v) mixture of epoxy embedding medium (Sigma, #45359-1EA-F) and ethanol for 1 hour, followed by a 66% (v/v) mixture of epoxy medium/ethanol for 2 hours, and pure epoxy medium overnight at 4 °C. Epoxy resin-filled TAAB capsules were placed upside down on the glass bottom of the imaging dish and cured at 60 °C for 48 hours. The glass bottom was removed by alternating submersion in boiling water and liquid nitrogen. The surface of the epoxy block was trimmed with a razor blade to a square at the previously noted localization coordinates of the acquired cell. Ultrathin serial sections of 300 nm for tomography were cut with an ultramicrotome (Leica Microsystems, UC6) equipped with a diamond knife (Science Services # DU3530). The ultrathin serial sections were transferred to a single slot grid (TEM, grid, 2×1 mm, slit, Cu, Science Services, #G2010-Cu) previously coated with 1% Pioloform® (Ted Pella, # 19244) in Chloroform (VWR, # 83626.290) and incubated for 10 min with protein A gold 10 nm (CMC Utrecht, #A-1904) diluted 1:150 in ddH2O as fiducials and washed 5 times in ddH2O. Sections were stained with 1.5% uranyl acetate (Agar Scientific, #R1260A) in ddH2O for 15 minutes at 37 °C and 3% Reynold’s lead citrate solution prepared from lead(II) nitrate (Roth, #HN32.1) and trisodium citrate dehydrate (Roth #4088.3) in ddH2O for 3 min. Grids were stored in grid boxes (Science Services, G71135-01) until imaging.

#### Electron tomography and CLEM image generation

Electron microscopy images were acquired using a JEM-2100 Plus transmission electron microscope (JEOL) operating at 200 kV for tomography equipped with a OneView 4K camera (Gatan). Transmission electron microscopy was used to identify the cells previously imaged by light microscopy based on the cell outline and shape of the mitochondrial network. Tomograms of 300 nm thick sections were then generated by tilting the sample along one axis from −60° to 60° using SerialEM (Mastronarde, 2005) and reconstructed using the Etomo plugin for IMOD (Table 1, ^53^). Overlays of TEM and STED images were generated using the EC-CLEM plugin of the image analysis software ICY (Table 1, ^54^). Since the 2D STED z-stacks have an axial resolution of less than 300 nm, a single z-plane of the 2D STED z-stack was aligned and overlayed with the correlating tomogram.

#### 3D rendering of tomograms

The MOM, MIM, *cristae*, and ER membranes were traced using the Microscopy Image Browser ver. 2.83 (MIB, Table 1, http://mib.helsinki.fi) with a line width of 3 to 4 pixels. Each membrane was redrawn separately. Positions with insufficient contrast in ER and *cristae* membranes were redrawn if the z-image above and below showed a continues membrane 3D rendering and visualization of the traced membranes was performed using Imaris software version 10.0.01 (Oxford Instruments, Table 1).

#### Correlation of BAK stoichiometry and size of BAK structures

The length of individual BAK line and arc structures as well as the perimeter of ring structures were measured in STED images using the segmented line tracing tool in ImageJ/FiJi (Table 1, ^51^). For stoichiometry/size correlation of individual BAK structures, line and arc lengths and ring perimeters were manually correlated with BAK stoichiometry derived from stoichiometric quantification of the photon-counting confocal image of the same cell. The correlation between stoichiometry and size was tested by calculating the non-parametric Spearman correlation coefficient r and the two-tailed P value at a 95% confidence interval.

#### Measurement of the size of MOM discontinuities

The size of MOM openings was measured using the line plot length tool in ImageJ/FiJi (Table 1, ^51^). The position of BAK foci was localized in the CLEM image, and the position with the highest contrast of the MOM in the tomogram was used to define the start and end points of the line plot. The fluorescence intensity of HALO-BAK and the opening of the MOM from the EM tomogram (Figure 1D-E) was measured using a (curved) line profile along the MOM opening with a width of 5 pixels. The fluorescence intensity was plotted using the line plot tool, from which the linear distance between signal maxima (BAK fluorescence) or minima (EM signal) was measured. We cannot discard that sample preparation for EM may affect the integrity of the MOM, which we did not detect as an issue by visual inspection of our samples. Yet, we specifically focused on MOM discontinuities associated with HALO-BAK assemblies, found as few discrete foci on mitochondria, further decreasing the likelihood that the MOM discontinuities analyzed could be artifactual.

#### MIM extrusion measurement and mitochondrial matrix swelling

To measure the extent of MIM extrusion through apoptotic pores in the presence or absence of osmotic mitochondrial matrix swelling, U2OS HALO-BAK cells were seeded in a glass-bottom 8-well μ-slide (IBIDI) at a density of 2×10^4^ cells per well one day before the experiment. Cells were stained with 300 nM Janelia Fluor X 650 HaloTag® Ligand for 1 hour at 37 °C to label BAK and with 150 nM MitoTracker™ Orange CMTMRos for 20 minutes at 37 °C to visualize the mitochondrial matrix. After labeling, the cells were washed twice with DMEM and incubated for 15 - 30 minutes at 37 °C to remove unbound dye. Apoptosis was induced using 10 µM ABT737, 10 µM S63845, and 10 µM Q-VD-OPh for 60 minutes in phenol red free DMEM at 37 °C and 5% CO_2_. For osmotic mitochondrial matrix swelling, cells were treated with 10 nM valinomycin (Sigma, #V0627) and 10 nM nigericin (Sigma, #N7143) in DMEM for 5 minutes. The cells were fixed in 4% (v/v) PFA in PBS for 10 minutes and incubated in PBS with 50 mM NH_4_Cl for 15 minutes at room temperature to quench unreacted fixative. The cells were permeabilized in PBS with 0.25% (v/v) Triton™ X-100 for 5 minutes, washed three times for 5 minutes in PBS, and blocked for PBS with 2% (w/v) BSA for 60 minutes at room temperature. The MOM was labeled with a 1:200 (v/v) dilution of rabbit α-TOM20 (Sigma, #HPA011562) in 2% (w/v) BSA overnight at 4 °C. After extensive washing with PBS, the cells were incubated with a 1:150 (v/v) dilution of Abberior STAR GREEN-coupled goat α-rabbit IgG secondary antibody (Abberior, #STGREEN-1002-500UG) for 1 hour at room temperature, followed by washing with PBS.

Alternatively, cells were seeded two days prior to the experiment (as described above) and transfected one day before the experiment with 50 ng TOM20-SNAP using Lipofectamine™ 2000. The cells were stained with 300 nM Janelia Fluor X 650 SnapTag® Ligand (Janelia Materials) for 30 minutes at 37 °C to label TOM20. In parallel, the MIM was labeled with 600 nM PKFO and 10 µM verapamil at 37 °C for 2 hours. Apoptosis and osmotic swelling of the mitochondrial matrix were induced as described above. The cells were fixed with 2% glutaraldehyde (in fixation buffer (100 mM HEPES/KOH pH 7.4, 3 mM CaCl_2_, 2.5% (w/v) sucrose, in HPLC water) for 20 minutes at room temperature and washed thoroughly with PBS.

Confocal/2D STED microscopy images of single cells were acquired using an inverted laser scanning STED microscope (TCS SP8 STED 3x, Leica, Wetzlar) equipped with a HC PL APO 100x/1.40 Oil STED_white_ objective (Leica, Wetzlar) and a tunable white light laser source (Leica, Wetzlar). JFX650 was excited at a wavelength of 640 nm, MitoTracker™Orange CMTMRos and PKFO at 561 nm, and Abberior STAR GREEN at 488 nm. STED de-excitation of JFX650 and PKFO was performed at 775 nm. The fluorescence emission signal of JFX650, MitoTracker™Orange CMTMRos and Abberior STAR GREEN or JFX650 and PKFO was detected line sequentially using hybrid detectors with a gating of 0.5 - 6 ns (for the STED image) and 8-fold line averaging. Images were adjusted for brightness and contrast using InageJ/Fiji (Table 1) and, where mentioned, deconvolved using Huygens Professional version 19.04 (Scientific Volume Imaging, The Netherlands, Table 1, http://svi.nl).

MIM extrusion was quantified by eye as intact mitochondria, mitochondria with MOM discontinuity, and mitochondria with or without MIM extrusion after osmotic matrix swelling.

#### Line tension measurement

To measure the line tension of MIM, we adapted the protocol proposed by Baumgart et al ^36^ based on the theory of Jülicher and Lipowsky ^55^. This protocol consists of tracing experimental images of axisymmetric mitochondria in such a way that the equatorial plane is orthogonal to the plane defined by the mitochondrial pore. Defining this reference plane allows us to obtain the local radius *R* and the tangent angle ψ, as a function of the arc length *S* (Figure 4J-K). *R*(*S*) and ψ(*S*) allow us to calculate the Hamiltonian, H, which is conserved on every point of the system (Figure 4K-L). This Hamiltonian is defined by the bending, pressure, tension, and line energy along the arc length, and in the limit of low temperatures, the following equality holds:

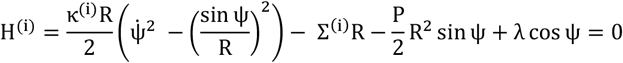

where κ^(i)^, Σ^(i)^, P, are the bending modulus, lateral tension, and outer excess pressure, respectively, of the mitochondria in the ith domain, i.e., MIM exposed to the cytosol and unperturbed MIM. The dot indicates differentiation with respect to the arc length. In this model, we do not take into account the energy contribution from the spontaneous curvature. *λ*(*S*) is a Lagrange multiplier that enforces the spatial constraint 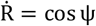. Its derivative with respect to *S* satisfies^55^:

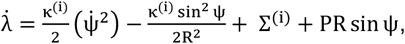

For the MIM inside (S < S^*^) and outside the MOM (S > S^*^), we interpret *λ*(S) as the radial force of the MOM onto the MIM, respectively. A step in *λ*(S) at the MOM pore then amounts to an effective line tension on the pore,

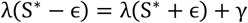

akin to the step in *λ*(S) at the domain boundary of the phase-separated system studied by Baumgart et al ^36^. The step height *γ* is the line tension to be determined, and *ϵ* an infinitesimal distance on either side of the boundary. *λ*(S) was obtained by integrating along the entire arc length and using the boundary condition, and used to calculate H(S) (Figure 4L).

Using *R*, ψ, and 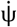 from the experimental images, we modulate the values of κ^(2)^, Σ^(i)^, P, and *γ* to minimize the sum of [H(S)]^2^. Here, we set κ^(1)^ = 15.48 KT as this value was reported for mammalian MIM in computational studies ^56^. As a validation of the above ansatz, we obtained a minimized value for the MOM of κ^(2)^ = 16.14 KT *≈* κ^(1)^, i.e., with the bending rigidity essentially unchanged as expected.

#### Modeling of MIM dynamics

The dynamics of MIM remodeling during BAX/BAK–mediated apoptosis was simulated using the TriMem code (Table 1) for a triangulated fluid-elastic model of the membrane ^57^. In TriMem, the position of the vertices, the topology of the connecting edges, and in turn the membrane shape evolve in time according to the so-called Helfrich Hamiltonian. Here, the dynamics of the MIM is subject in addition to its confinement by the MOM structure.

As a minimal model of the *cristae* structures in an intact mitochondrion, we placed a stack of 6 flat disk-shaped membranes inside a tubular membrane lining the MOM container. By establishing additional edges, we created necks to fuse the disk shapes with the tubular membrane. The resulting mesh contained 11399 vertices and 22794 triangular faces. To set up, smoothen, and subdivide the initial mesh structures, we used the Autodesk Fusion360 software (Table 1).

#### Modelling of MOM and its interaction with the MIM

We described the MOM of an intact mitochondrion as a rigid pill-shaped container confining the MIM (Figure S5). The container consists of a cylinder of radius **R**_**container**_ = **0. 71 μm** and height **h**_**container**_ = **0. 74 μm** aligned with the **Z** axis that is capped at its ends by outward-facing hemispheres of radius **R**_**container**_. To model the confinement of the MIM in the MOM, we added a repulsive potential on the vertices of the MIM mesh as a function of their normal distance **r** from the container,

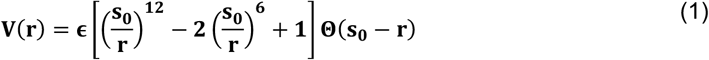

where **Θ** is the Heaviside function, **s**_**0**_ is the length scale of the confining potential, and **ϵ** its characteristic energy whose values, as listed in Table S1.

To describe the BAX/BAK-mediated formation of a pore in the MOM, we introduced a circular opening in the external container. Following the observations in the microscopy experiments, we placed the pore center axially at **z** *>* **0, x** = **y** = **0** at one end of the mitochondrion. With a polar angle of ‘ALPHA (**α**)’ on the hemispherical cap quantifying the size of the opening (SI), the radius of the pore is **r**_**pore**_ = **R**_**container**_ **sin**(**α**). For MIM vertices **i** at Cartesian position (**x**_**i**_, **y**_**i**_, **z**_**i**_), we similarly defined a polar angle 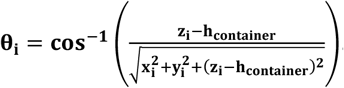. For **θ**_**i**_ ≥ **α**, we defined the distance **r** in Eq. (1) as the closest distance to the cylinder and the two hemispheres; for **θ**_**i**_ *<* **α**, we instead set **r** to the minimum distance **Δr**_**i**_ to the circular rim,

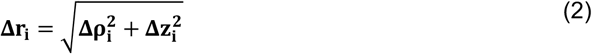

where **Δρ**_**i**_ and **Δz**_**i**_ denote the shortest radial and vertical distances between the rim and the vertex **i** in cylindrical coordinates,

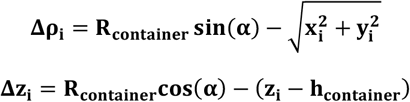

We define the contributions of the latter terms to the potential energy as the rim potential. From the associated forces exerted by the MIM on the pore rim, we deduced an effective line tension, as explained below.

#### Simulations

The initial MIM shape mimicking an intact mitochondrion was equilibrated with TriMem (Table 1). The hybrid Monte Carlo production runs were performed using the parameters provided in SI. The integration time step in the Hamiltonian dynamics segments of the hybrid Monte Carlo (hMC) runs was set to **1** × **10**^**−4**^**τ**, where **τ** is the reduced time unit. Each segment consisted of 50 time steps, thus lasting **0. 005τ**. The simulations were run at reduced temperature **T** = **1** and vertex mass **M** = **0. 2** with bending rigidities **κ**_**B**_ = **10 k**_**B**_**T** ^38^ and **60 k**_**B**_**T**, respectively. Stiffening the membrane tended to speed up the membrane evagination. In separate runs, we examined the effect of an osmotic inflation of a leaky MIM by ramping up the matrix volume after time point **7500τ**. For this, we added a term 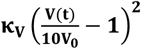 to the potential energy (eq 7 in ^57^), where **V**_**0**_ is the initial volume of the matrix and **V**(**t**) the instantaneous volume. Starting from zero, we increased the coupling constant **κ**_**V**_ every **500τ** by **5000k**_**B**_**T**.

We performed simulations with varying pore opening angles of **α** = **0. 25, 0. 5, 0. 75, 1. 0**, and **1. 5** radians, which correspond to reduced pore radii of 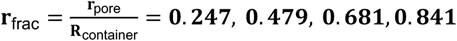 and **0. 997**. Note that the MOM shape and pore opening are fixed in each replica run.

#### Pore line tension evaluation

The MIM evaginating through the pore exerts forces onto the rim of the pore in the MOM. We convert these forces into an effective line tension (**γ**) decomposed into two contributions: (1) 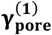 accounting for forces associated with direct vertex-rim interactions, as captured by the rim potential and (2) 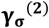 accounting for pore expansion driven by tension in the MOM. We estimate the former as the mean outward directed force on the rim in plane with the pore,

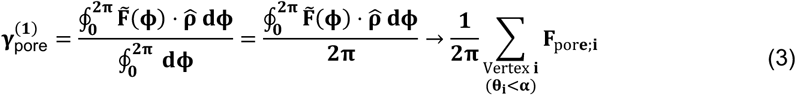

where **ϕ** is the angle in the cylindrical coordinate system, 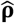 is a radial unit vector normal to the **Z** axis, and 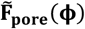 is the force exerted on the pore rim by the MIM. In the triangulated membrane model, **F**_**pore;i**_ denotes the radially directed component of the force a vertex **i** exerts on the pore rim, which we calculate from the gradient of the container potential **V**(**r**) as

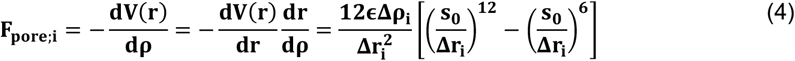

where **Δr** and **Δρ** denote the closest distance and the closest lateral distance of the vertex **i** from the pore rim as defined in Eq. (2).

In addition to the direct interaction between the MIM and the pore rim inducing 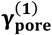, we account for an expansive force induced by the MIM pushing up against the MOM for **θ**_**i**_ ≥ **α**. These forces normal to the MOM create an effective excess pressure (**ΔP**) acting on the MOM. As a curved surface, this pressure in turn induces a surface tension ***σ*** in the MOM. For a sphere of radius **R**_container_, this tension is given by the Laplace-Young relation, 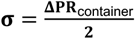, which we use here as a rough approximation also for the model mitochondria. We calculate the average pressure gradient across the MOM as

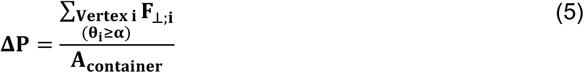

where 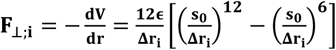 is the normal force on the container exerted by vertex I for **r** *<* **s**_**0**_ outside the pore angle **α**. 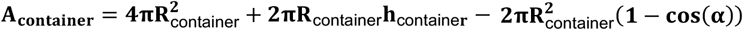 denotes the area of the container, with the spherical cap cut out by the pore subtracted.

We convert the tension ***σ*** in the MOM into a line tension in the pore by requiring mechanical stability,

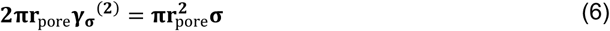

adopting the relation for a planar membrane. The resulting effective line tension is then

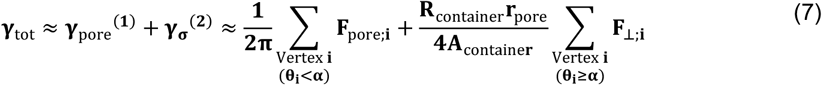

We evaluate this expression for the structures saved along the TriMem simulations to obtain an effective line tension on the BAX/BAK-pore as a function of time, as the MIM evaginates through the pore. With physical units for the container dimensions and membrane bending rigidity, the effective line tensions has units of force (i.e., pN).

#### Data representation and statistical analysis

All correlative stoichiometry super-CLEM data are representative or individual examples of n = 8 independent experiments (Figures 1, 2, S2 and S3), except for the classification of MOM pore categories (Figures 3 and 5C-G), which represents n = 6 independent experiments. All other data are representative of at least n = 3 independent experiments (Figures 4 and S1), as indicated in the respective figure legends.

Quantitative data (in Figures 2B-C, and J, 4C, H and M, 5G, and S2F-G) were tested for normality using the Shapiro-Wilk normality test. Data with a normal distribution are presented as mean ± SD, while data that do not follow a normal distribution are presented as median ± interquartile range (IQR), both together with the individual data points. Data containing at least one non-normally distributed data set along with otherwise normally distributed data sets were treated as non-normally distributed.

Significance levels of non-normally distributed data and data with more than two unpaired data sets were assessed using non-parametric one-way ANOVA (Kruskal-Wallis test) and Dunn’s multiple comparison test to determine multiplicity adjusted P values (family-wise significance and confidence level set to 0.05, Figures 2B-C and J, 4C, and 5G). Normally distributed data with two unpaired data sets were tested for significance using unpaired non-parametric Student’s t-test (Mann-Whitney test, Figure 4H). Corresponding P values are reported in the respective figures.

## Data availability

This study includes no data deposited in external repositories.

## Author contributions

A.J. and T.D. implemented CLOSE microscopy and performed research, with support from F.G., K.S. and A.S. C.Z. generated and characterized the HALO-BAK cell line. D.G. perfomed line tension analysis from images. H.K. and G.H. set up and analyzed the membrane dynamics simulations with help from J.K. A.G.S. conceived the project and supervised research. A.J. and A.G.S. wrote the manuscript with the help of all other authors.

## Acknowledgements

We thank the CECAD imaging facility for excellent assistance and specifically Dr. Christian Jüngst for his outstanding technical and methodological support regarding image acquisition and analysis for stoichiometry quantification and STED microscopy. We further thank Gudrun Zimmer for technical support. This work was funded by the European Research Council (ERC-Co 817758) and the Deutsche Forschungsgemeinschaft (DFG, German Research Foundation) – 654651/GRK2364 MOMbrane, DFG-INST 2016/742-1 FUGB and DFG-INST 216/793-1 FUGG, and the Max Planck Society. We thank the Max Planck Computing and Data Facility (MPCDF) for computational support and Sebastian Kehl for help with the TriMem membrane dynamics simulation code. H.K thanks International Max Planck Research Schools on cellular biophysics (IMPRS-CBP).

## Declaration of AI and AI-assisted technologies in the writing process

During the preparation of this work the authors used DeepL Write (DeepL SE) in order to improve readability and language of the text. After using this tool, the authors reviewed and edited the content as needed and take full responsibility for the content of the publication.

