## Supplementary information for "Mechanical forces drive mitochondrial matrix extrusion and apoptotic pore growth"

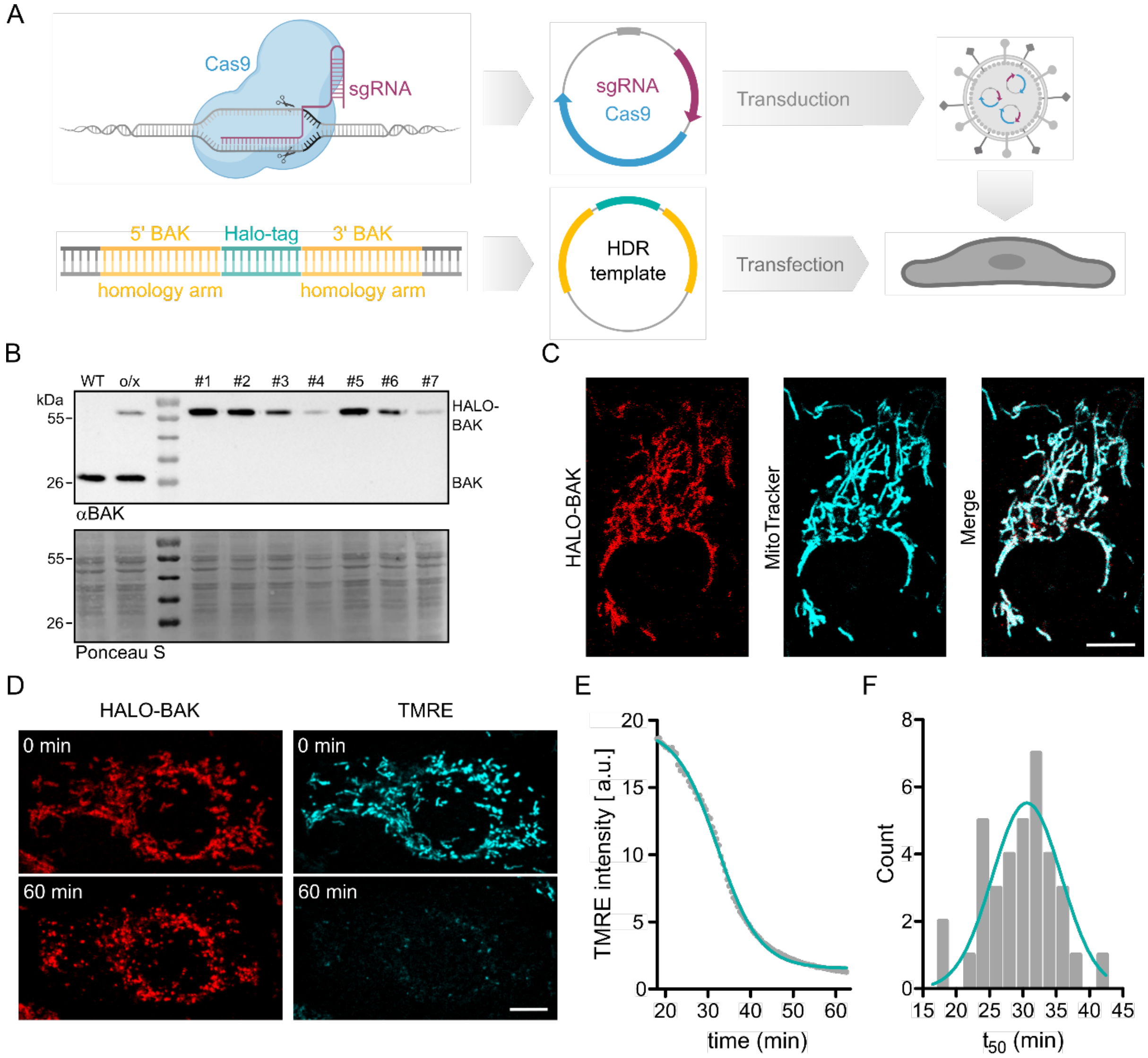


**Figure S1: Generation and validation of the U2OS HALO-BAK cell line.** A) We generated a U2OS cell line expressing BAK genomically tagged with HALO-tag (HALO-BAK) using CRISPR/Cas9, which allows visualization of BAK with advanced organic fluorophores at endogenous expression levels. A) shows the schematic representation of the workflow using a combined CRISPR/Cas9 approach: A synthetic guide RNA (sgRNA) specific for the endogenous locus of BAK is cloned into a lentiviral expression vector together with the coding sequence of the Cas9 enzyme (upper panel). Homology arms of 800 bp of the genomic BAK locus upstream (5' BAK homology arm) and downstream (3' BAK homology arm) of the start codon are fused to the coding sequence of the HALO tag and cloned into a homology-directed repair (HDR) template expression vector (lower panel). Both the Cas9/sgRNA and the HDR template are delivered to the target cell by lentiviral transduction or transfection, respectively. B) In addition to PCR genotyping (not shown), we performed immunoblotting of single cell clones to confirm homozygous genome editing. B) shows the expression of the HALO-BAK fusion protein in single cell clones (#1-7) by immunoblotting against BAK. Cell lysates from wild-type U2OS cells (WT) and wild-type cells transiently overexpressing HALO-BAK (o/x) were used as controls. Whole protein staining (Ponceau S) is shown to control for equal sample loading. C-F) Live-cell confocal microscopy of HALO-BAK in healthy cells and during apoptosis demonstrates the functionality of genome-edited HALO-BAK, which presented the same subcellular localization and comparable cell death kinetics as described for wild type BAK. C) Representative confocal fluorescence microscopy image demonstrating the subcellular localization of HALO-BAK. HALO-BAK was labeled with JFX650 HALO-Tag ligand (red) and mitochondria were visualized using MitoTracker green (cyan) as a reference. Scale bar 10 µm. D) Representative confocal fluorescence microscopy images of HALO-BAK (red) before (0 min) and after induction of cell death (60 min). Mitochondria were stained with the potential-sensitive dye TMRE (cyan) to monitor the loss of mitochondrial membrane potential over time. Scale bar 10 µm. E) Representative quantification of TMRE signal intensity over time. The data were fitted with an exponential decay function (blue) to extract the time of 50% TMRE signal loss ($t_{50}$). F) Distribution of $t_{50}$ values fitted with a Gaussian function (blue) to extract the mean time of 50% TMRE signal loss. Experiments are representative of n = 3 independent experiments with n = 20 cells each.

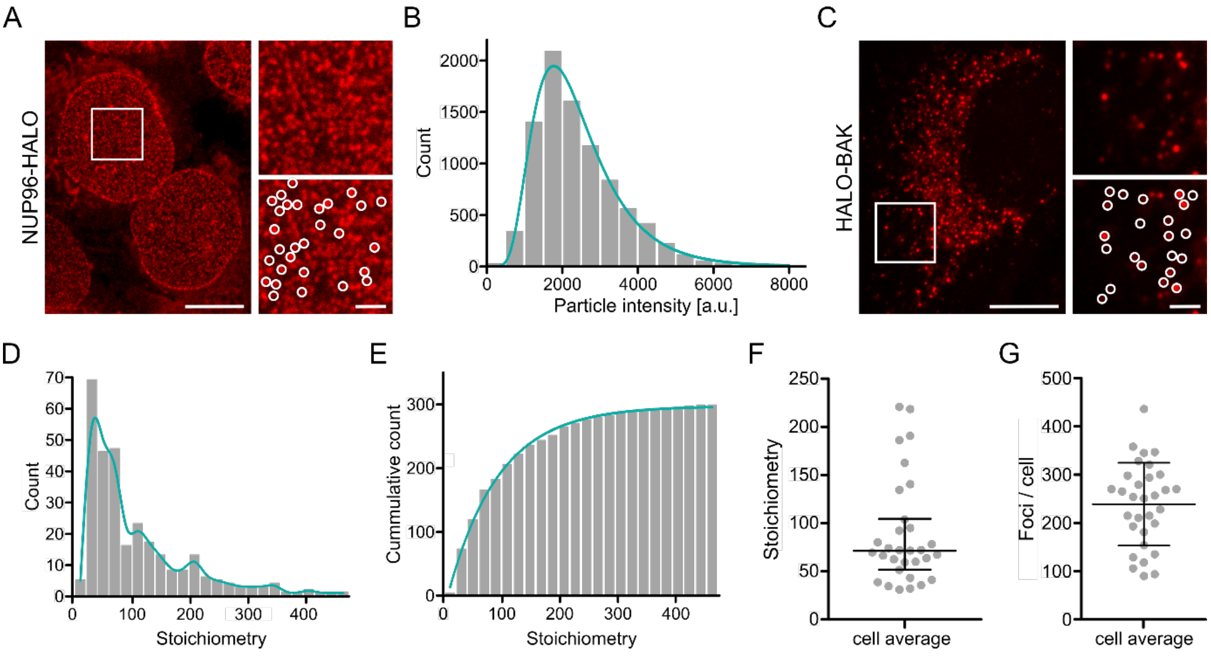


**Figure S2: Stoichiometric quantification of HALO-BAK stoichiometry.** We performed single particle brightness quantification of individual NUP96-HALO complexes (A-B), which allows ratiometric calculation of the theoretical brightness of a monomeric fluorescent emitter, which we used to convert the brightness of HALO-BAK apoptotic particles to HALO-BAK molecularity (C-E). A, C) Representative photon-counting confocal microscopy image of U2OS NUP96-HALO (A) and U2OS HALO-BAK (C) labeled with JFX650 HALO-Tag ligand. With the exception of apoptosis induction, U2OS NUP96-HALO cells were prepared and imaged with photon-counting confocal microscopy in the same manner as U2OS HALO-BAK samples. Right panels represent zoomed regions as indicated by the rectangle. Detected regions of interest of measured HALO-BAK and NUP96-HALO foci, respectively, are shown in the lower right panels (white circles). Scale bar 10 µm, zoomed images 2 µm. Images are representative of n = 123 NUP96-HALO and n = 31 HALO-BAK cells from n = 8 independent experiments. B) Exemplary fluorescence intensity distribution of n = 1937 individual NUP96-HALO foci measured from n = 22 individual cells of one measurement day. Lognormal fitting was applied to extract the median particle intensity of the population (blue line), which is used as the calibration standard for radiometric stoichiometry quantification. D) Population distribution (gray) and probability density function (blue line) of the stoichiometry of HALO-BAK foci detected from the cell shown in (C). E) Cumulative distribution (gray) of HALO-BAK stoichiometry in (D) fitted with an exponential decay function (blue line) to extract the average stoichiometry of HALO-BAK foci in the given cell. F, G) Quantification of median cellular BAK stoichiometry in apoptotic mitochondria (F) and number of HALO-BAK foci detected per cell (G). Values are presented for individual cells (individual data points) as well as the median (F) or mean (G, line) ± interquartile range (F) or SD (G, whiskers) of n = 31 cells from n = 8 independent experiments.

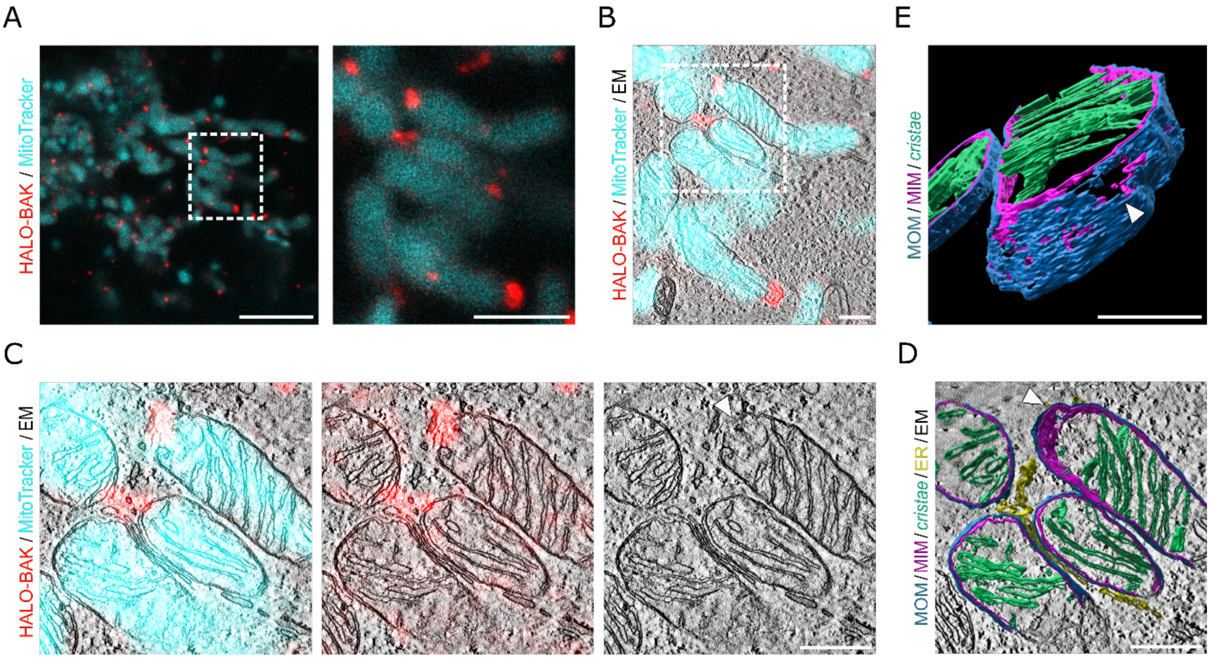


**Figure S3: Correlative suplerCLEM analysis of a single mitochondrion.** A) Representative STED microscopy image of U2OS HALO-BAK cells labeled with JFX650 HALO-Tag ligand (red) one hour after apoptosis induction. Mitochondria were labeled with MitoTracker orange (cyan, confocal). Enlarged image (right panel) corresponds to the cropped region as indicated. Scale bar 5 µm, magnified image 2 µm. B) Overlaid STED-CLEM image of a single mitochondrion showing MitoTracker (cyan, confocal) and BAK (red, STED) fluorescence signal and EM image (grayscale). The dashed rectangle indicates the cropped region shown in (C-D). Scale bar 500 nm. C) Enlarged image of cropped region from image in (B) showing MitoTracker (cyan, confocal) and BAK (red, STED) fluorescence signal and EM image (grayscale, left panel), BAK signal and EM image (middle panel), and single EM image (right panel). Scale bar 500 nm. D) 3D rendered MOM (blue), MIM (purple), *cristae* (green), and ER membranes (yellow) overlaid with EM image (grayscale) in top view. Scale bar 500 nm. Scale bar 500 nm. E) Tilted and magnified view of 3D rendered MOM (blue), MIM (purple), and *cristae* membranes (green). Scale bar 250 nm. Arrowheads in (C-E) indicate position of MOM and MIM discontinuities.

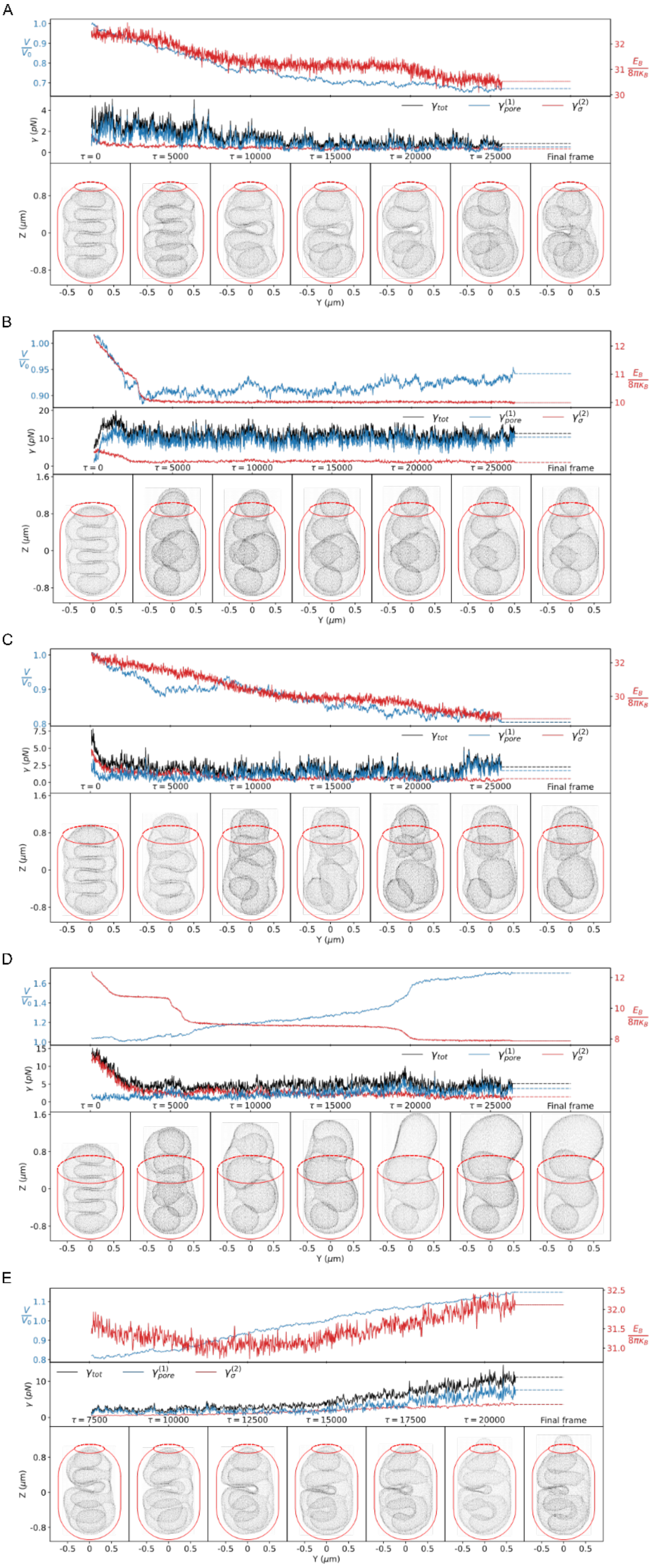


**Figure S4: Membrane dynamic simulation of MIM extrusion.** A-D) MIM (grey in bottom panel) contained in MOM (red schematic outline) with a pore of size $\alpha = 0.50$ corresponding to a relative diameter of $r_{\mathrm{frac}} = 0.479$ and a bending rigidity of $\kappa_B = 10\,k_B T$ (A), $\alpha =$

$0.75; r_{\text{frac}} = 0.682; \kappa_B = 60 k_B T$ (B), $\alpha = 1.00; r_{\text{frac}} = 0.841$; $\kappa_B = 10 k_B T$ (C), and $\alpha = 1.50; r_{\text{frac}} = 0.997$; $\kappa_B = 60 k_B T$ (D) simulated without osmotic shock. E) A pore of size $\alpha = 0.50$ corresponding to a relative diameter of $r_{\text{frac}} = 0.479$ and a bending rigidity of $\kappa_B = 10\ k_B T$ with osmotic inflation starting from time point $\tau = 7500$. The bottom panels show snapshots at the indicated time points, the top panels the membrane bending energy $E_B$ (red) and the volume relative to that of the starting structure (blue). The center panel shows the effective line tension created by the evaginating membrane in the BAX/BAK-pore (see Methods for components).

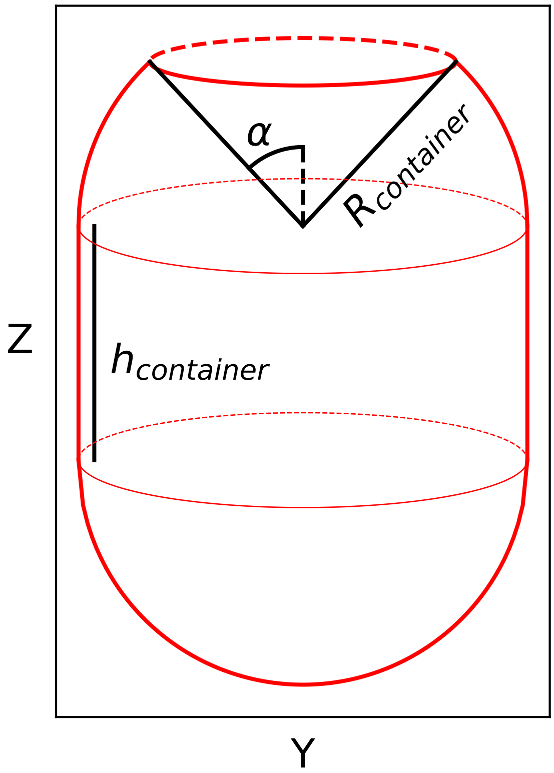


**Figure S5: Schematic diagram of the rigid container mimicking the MOM.** The container consists of a cylindrical central body of radius $\mathbf{R_{container}}$ and height $\mathbf{h_{container}}$ aligned with the $\mathbf{Z}$ axis that is capped at its ends by hemispheres of radius $\mathbf{R_{container}}$. The circular opening along the positive $\mathbf{Z}$ axis describes the apoptotic pore on the MOM. The size of the pore is defined by $\boldsymbol{\alpha}$, being the polar angle from the positive $\mathbf{Z}$ axis to the rim of the pore, taking the center of the intact hemispherical cap as the reference point. The pore radius is then $\mathbf{r_{pore}} = \mathbf{R_{container}} \, \mathbf{sin}(\boldsymbol{\alpha})$.

**Table S1**

[GENERAL]

| | |
|---|---|
| algorithm | hmc |
| info | 10 |
| input | IMM.stl |
| output_prefix | out/out |
| restart_prefix | out/out |
| checkpoint_every | 1000 |
| output_format | vtu |

[HMC]

| | |
|---|---|
| num_steps | 10000000 |
| step_size | 1e-4 |
| traj_steps | 50 |
| momentum_variance | 0.2 |
| thin | 500 |
| flip_ratio | 0.025 |
| flip_type | parallel |
| initial_temperature | 1 |
| cooling_factor | 0 |
| start_cooling | 10000000 |

[BONDS]

| | |
|---|---|
| bond_type | Edge |
| r | 1 |

[SURFACEREPULSION]

| | |
|---|---|
| n_search | cell-list |
| rlist | 0.5 |
| exclusion_level | 0 |
| refresh | 1 |
| lc1 | 0.155 |

| | |
|---|---|
| r | 1 |

[EXTERNALPOTENTIAL]

| | |
|---|---|
| type | porous_MITO |
| epsilon | 1 |
| sigma | 0.5 |
| radius | 3.55 |
| half_height | 1.85 |
| alpha | 0.25/0.50/0.75/1.00/1.50 |

[ENERGY]

| | |
|---|---|
| kappa_b | 10/60 |
| kappa_a | 1000 |
| kappa_v | 0/$\boldsymbol{\kappa_V(\tau)}$ |
| kappa_c | 0 |
| kappa_t | 1000 |
| kappa_r | 1000 |
| kappa_e | 1 |
| area_fraction | 1.0 |
| volume_fraction | 10.0 |
| curvature_fraction | 0.01 |
| continuation_delta | 0.0 |
| continuation_lambda | 1.0 |

**Table S1: List of parameters used in the TriMem simulations.** The desired algorithm and in/output formats are defined in [GENERAL]. As listed in [HMC], the integration time step in the Hamiltonian dynamics segments of the hybrid Monte Carlo runs was set to $\mathbf{1 \times 10^{-4}\tau}$, where $\boldsymbol{\tau}$ is the reduced time unit. The duration of the each segment was 50 time steps, hence, $\mathbf{0.005\tau}$; an output vtu file is written every $\mathbf{2.5\tau}$. The simulations were run at reduced temperature $\mathbf{T = 1}$ and mass $\mathbf{M = 0.2}$. The simulations were halted soon after $\mathbf{25000\tau}$ for the replicas without the volume constraint, and around $\mathbf{20000\tau}$ for the osmotic shock cases.
[EXTERNALPOTENTIAL] defines the type and shape of the external container, where epsilon $(\epsilon)$, sigma $\left(\sigma = 2^{-\frac{1}{6}}s_0\right)$, radius $(R_{container})$, half_height $(0.5h_{container})$ and alpha $(\alpha)$ denote characteristic energy scale, length scale of the confining potential, radius and half the height of the cylindrical central body of the container mimicking the OMM in internal units, and the pore size, respectively. [ENERGY] defines the energy penalty constants of the Helfrich

Hamiltonian [1], and the corresponding reference properties of the IMM. Note one set of the simulations was conducted without the volume penalty $(\kappa_V = 0)$, while another set of replica simulations was continued from $7500\tau$ with the volume constraint, in which $\kappa_V$ was increased by $5000 k_B T$ every $500\tau$ with a target of $10V_0$.


1. Siggel, M., Kehl, S., Reuter, K., Köfinger, J. & Hummer, G. TriMem: A parallelized hybrid Monte Carlo software for efficient simulations of lipid membranes. *J. Chem. Phys.* **157**, 174801 (2022).
